# Integrative analysis of the genome, transcriptome, and proteome identifies causal mechanisms of complex traits

**DOI:** 10.1101/2024.03.28.587202

**Authors:** Jeffrey Okamoto, Xianyong Yin, Brady Ryan, Joshua Chiou, Francesca Luca, Roger Pique-Regi, Hae Kyung Im, Jean Morrison, Charles Burant, Eric B. Fauman, Markku Laakso, Michael Boehnke, Xiaoquan Wen

**Affiliations:** Department of Biostatistics and Center for Statistical Genetics, University of Michigan, Ann Arbor, MI 48109, USA; Department of Epidemiology, School of Public Health, Nanjing Medical University, Nanjing, Jiangsu 211166, China; Internal Medicine Research Unit, Pfizer Worldwide Research, Development and Medical, Cambridge, MA 02139, USA; Center for Molecular Medicine and Genetics, Wayne State University, Detroit, MI 48201, USA; Section of Genetic Medicine, Department of Medicine, University of Chicago, Chicago, IL 60637, USA; Department of Internal Medicine, University of Michigan, Ann Arbor, MI 48109, USA; Institute of Clinical Medicine, Internal Medicine, University of Eastern Finland and Kuopio University Hospital, Kuopio 70210, Finland

## Abstract

We present multi-integration of transcriptome-wide association studies and colocalization (Multi-INTACT), an algorithm that models multiple gene products (e.g. encoded RNA transcript and protein levels) to implicate causal genes and relevant gene products. In simulations, Multi-INTACT achieves higher power than existing methods, maintains calibrated false discovery rates, and detects the true causal gene product(s). We apply Multi-INTACT to GWAS on 1,408 metabolites, integrating the GTEx expression and UK Biobank protein QTL datasets. Multi-INTACT infers 52% to 109% more metabolite causal genes than protein-alone or expression-alone analyses and indicates both gene products are relevant for most gene nominations.

## 1 Introduction

Genome-wide association studies (GWAS) have greatly advanced our understanding of the genetic basis of complex diseases and traits by identifying numerous variant-level genetic associations. However, most identified associated variants lie in noncoding genome regions [1, 2], often obscuring their target genes and complicating the therapeutic target identification. Recent efforts to address this problem have resulted in the emergence of genome-scale molecular quantitative trait loci (QTL) annotations [3, 4] and the development of statistical methods bridging genetic variants with molecular phenotypes of gene candidates and complex traits [5, 6, 7, 8, 9, 10, 11, 12, 13, 14, 15, 16, 17, 18]. These mechanism-aware putative causal gene (PCG) implication methods [8] focus on molecular phenotypes that can be unambiguously linked to specific genes. Henceforth, we refer to such molecular phenotypes as *gene products*. Key gene products commonly used in QTL mapping include transcriptome abundance, isoform usage, RNA decay rate, and protein abundance.

Existing mechanism-aware methods primarily implicate PCGs through a single gene product. Transcriptome-wide association studies (TWAS) [10, 11, 12, 14, 15] evaluate gene expression’s mediating role by examining the correlation between genetically-predicted expression with a GWAS trait. Colocalization analyses [14, 16, 17] aim to identify variant-level overlap of causal expression QTLs (eQTLs) and GWAS associations. Each of these methods has been shown to have distinct limitations due to statistical and biological factors such as horizontal pleiotropy and linkage disequilibrium (LD) hitchhiking, which result in false positives and negatives [5, 19, 20].

While eQTL data have shown promise for expanding our understanding of complex trait genetic architecture [3, 21, 22], they do not always illuminate the true effects of causal genes on complex traits [23, 24]. Recent work [25] suggests that most disease heritability cannot be explained by tissue-specific *cis*-expression quantitative trait loci (eQTL) data but rather by other molecular mechanisms. In particular, splicing QTLs (sQTLs) [26, 3] and protein QTLs (pQTLs) [24, 27] have shown to display minimal eQTL overlap and may independently influence disease heritability. Consequently, transcriptome-wide association studies (TWAS) and colocalization analysis of GWAS and eQTL data are often ineffective means of identifying causal genes. Numerous molecular phenotypes – including isoform usage, RNA degradation, protein abundance – have demonstrated their relevance in explaining and predicting disease risk [28, 29, 30, 31, 32, 33, 34, 35, 36, 37, 38, 39, 40, 41]. Joint analysis of multiple gene products can offer a holistic view of the underlying biology and improve the PCG implication performance.

In this study, we introduce Multi-INTACT, a mechanism-aware PCG inference method that aggregates colocalization and TWAS evidence across diverse gene products within a Bayesian framework. Multi-INTACT gauges the causal significance of a target gene concerning a complex trait and identifies the pivotal gene products. Using comprehensive simulations, we demonstrate the advantages of Multi-INTACT over existing methods. Finally, we use Multi-INTACT to detect PCGs influencing plasma metabolite levels via RNA transcript or protein levels.

## 2 Results

### 2.1 Method Overview

The key idea of Multi-INTACT is to leverage information from *multiple* molecular phenotypes to implicate PCGs. Particularly, we define *gene products* as the molecular phenotypes that can be explicitly linked to genes. Contemporary experimental technology allows for the measurement of various gene products such as RNA abundance, isoform usage, RNA decay rate, and protein abundance. To examine potential gene-trait causative links, Multi-INTACT extends the canonical single-exposure (i.e., a single molecular phenotype) instrumental variables (IV) analysis/TWAS method to account for multiple endogenous variables, integrating colocalization evidence in the process. Genetic association analysis results of molecular QTLs and GWAS loci are essential to Multi-INTACT’s inferential procedure. We summarize the Multi-INTACT method workflow in Figure 1a.

**Figure 1:**
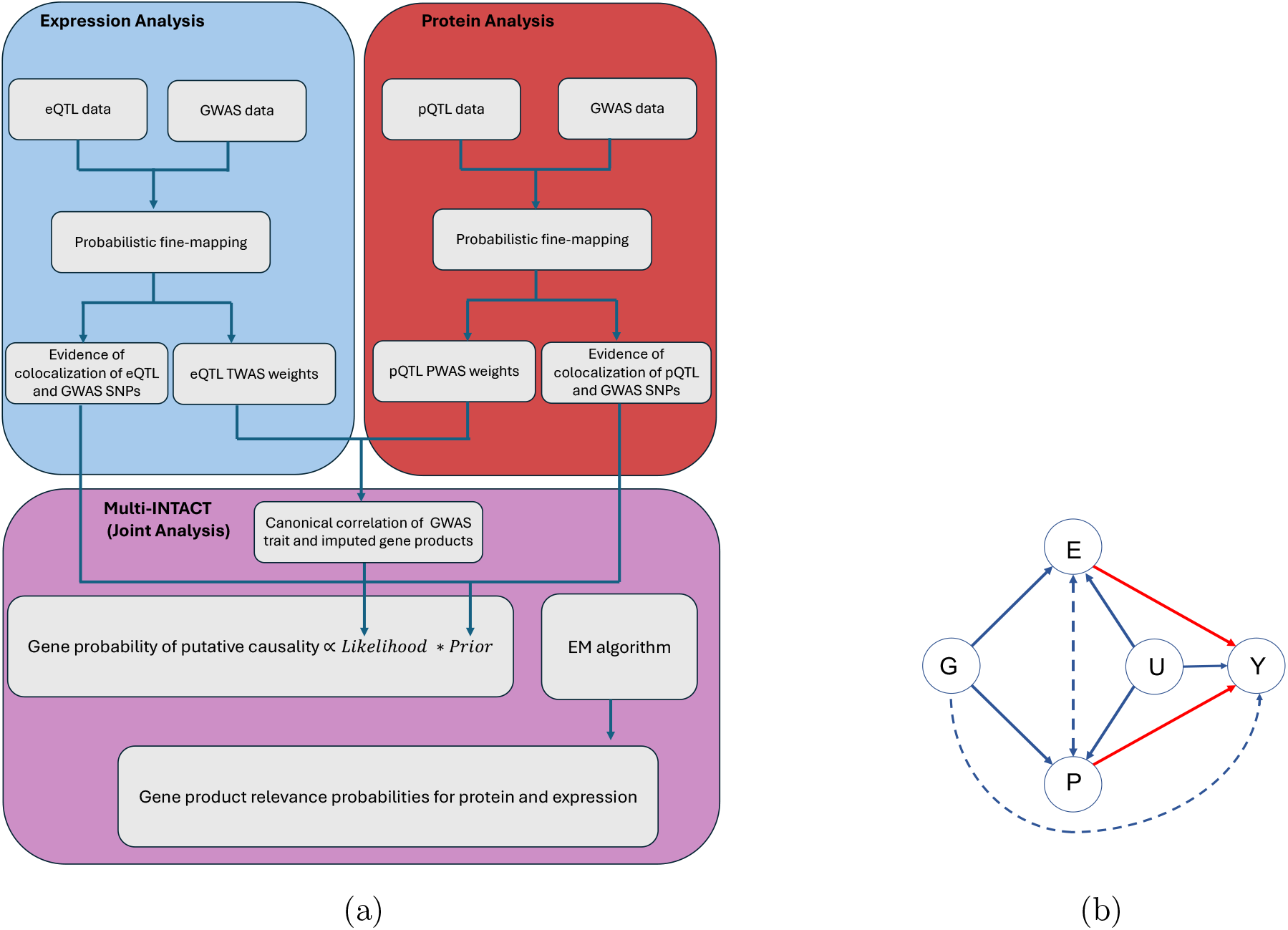
(a), Multi-INTACT workflow. (b), Causal diagram that connects genetic variants *G*, expression levels of a candidate gene *E*, protein levels of a candidate gene *P*, and a complex trait *Y* . The node *U* represents latent confounders of effects between *E*, *P*, and *Y* . A similar diagram is assumed by multivariable Mendelian Randomization methods. The edges that are highlighted in red are those that Multi-INTACT is designed to infer.

While the proposed inference framework can be applied to incorporate many gene products simultaneously, for simplicity, we illustrate the method using two gene products, gene expression levels (*E*) and protein abundance (*P*). The Multi-INTACT model is built upon the existing IV analysis framework, accommodating multiple endogenous variables (i.e., *E* and *P*) [42]. Figure 1b depicts the assumed directed acyclic graph (DAG) for potential relationships between molecular QTLs (*G*), gene products (*E* and *P*), unobserved confounding (*U*), and the complex trait of interest (*Y*). The primary goal of the statistical inference is to test for potential causal relationships from the gene products to the complex trait (i.e., *E → Y* and *P → Y*) represented by the do-calculus, *P* (*Y | do*(*E*)) and *P* (*Y | do*(*P*)) while allowing flexible relationships between gene products. More specifically, we aim to answer the following two related scientific questions for each gene candidate:

1. Is the gene candidate a PCG, i.e., do *any* of its gene products exert a potential causal effect on the trait of interest?
2. Provided that the gene candidate is a PCG, what are the relevant gene products?

To assess the plausibility of a gene candidate being a PCG, Multi-INTACT applies an empirical Bayes procedure to compute a gene-specific posterior probability, denoted as the *gene probability of putative causality*. In particular, the likelihood computation generalizes from the existing single-trait TWAS methodology by constructing the composite instrument variables, *E*^^^ and *P*^^^, using their respective molecular QTLs and subsequently testing their *canonical correlation* with *Y* . These composite IVs are naturally interpreted as the genetic prediction of the corresponding molecular phenotypes, and the procedure follows the principles of the IV analysis framework [43, 44]. The prior formulation is designed to ensure the key causal assumptions and validate the putative causality claim. Specifically, the prior incorporates the colocalization evidence of molecular QTLs and GWAS hits to guard against violations of the exclusion restriction (ER) assumption caused by widespread LD in genetic data. We note that under various possible causal scenarios (Figure S1), not all types of molecular QTLs are necessarily colocalized with GWAS hits. Additionally, considering the low practical power to detect colocalization [19], our implementation requires at least one type of molecular QTL to exhibit modest colocalization evidence as a minimum necessary condition for a candidate gene to be classified as a PCG. Finally, the complement of a gene probability of putative causality is a local false discovery rate (lfdr), which is suitable for the standard Bayesian false discovery rate (FDR) control procedure to guard against type I errors.

To address the second question on identifying relevant gene products, we formulate a model selection problem considering all possible causal relationships from examined gene products to the complex trait. In our illustrative example, Multi-INTACT evaluates the posterior probabilities for the following four mutually exclusive models:

1. *M*_0_: neither *E* or *P* exerts an effect on *Y*, i.e., the null model
2. *M_E_*: only *E* exerts an effect on *Y*, i.e., the *E → Y* model
3. *M_P_* : only *P* exerts an effect on *Y*, i.e., the *P → Y* model
4. *M_E_*_+_*_P_* : both *P* and *E* exert effects on *Y*, i.e., the (*E, P*) *→ Y* model

For a general case where there are *p* gene products considered, there are 2*^p^* models to be compared. This presents a practical limitation for Multi-INTACT, as the number of models increases rapidly with *p*.

Multi-INTACT implements an EM algorithm to estimate the empirical Bayes prior distribution among the three competing alternative models by pooling information across all gene candidates. It subsequently evaluates the posterior model probabilities, i.e., Pr(*M_E_ |* data), Pr(*M_P_ |* data), and Pr(*M_E_*_+_*_P_ |* data), for each gene candidate using the Bayes rule. Finally, Multi-INTACT reports a *gene product relevance probability* of each gene product for a given candidate by marginalizing the corresponding posterior model probabilities:

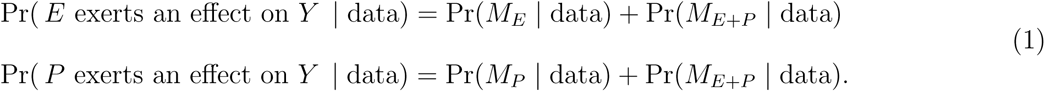

We provide technical details and features of the Multi-INTACT method in Methods. In brief, the Multi-INTACT model can be represented by a structural equation model (SEM) that allows pleiotropic effects. Hence, it is robust against some of the most common violations of the ER assumptions using genetic data. Importantly, although the causality claim is derived based on the one-sample design (i.e., all genetic, molecular phenotypes, and complex traits are measured on a single cohort), it can be extended to multi-sample designs, which are common among available genomic and genetic data.

### 2.2 Simulation Study

To evaluate the performance of Multi-INTACT, we perform extensive simulation studies based on genetic data from GTEx. We extend the simulation design introduced in [8], which uses real genotypes of 477K SNPs on chromosome 5 from 500 GTEx samples. The selected genomic region contains 1198 consecutive genes, each with at least 1500 common *cis*-SNPs, some of which are located in the overlapping cis-regions of multiple gene candidates. We consider a multi-sample design to simulate the molecular and GWAS phenotypes, where the residuals of *E, P*, and *Y* are uncorrelated after controlling for their shared genetic components. The phenotype data are generated by randomly sampling the DAGs shown in the second row of Table S1. Note that the data generative models differ from the Multi-INTACT model, as they make additional assumptions connecting *G, E,* and *P* . Each assembled simulated dataset contains 1198 genes with *∼* 80% non-PCGs and *∼* 20% PCGs. The causal mechanism of each simulated PCG follows a discrete distribution of *M_E_, M_P_*, and *M_P_* _+_*_E_* models, which is varied across different datasets. In total, we generate 100 datasets for analysis. Complete simulation details are provided in Methods and Supplemental Methods.

For each simulated dataset, we perform fine-mapping analyses using individual-level genotype-phenotype data for all molecular and complex traits. We then separately conduct colocalization and TWAS analysis for the protein-GWAS and expression-GWAS data. The resulting single-molecular trait integrative analysis data are subsequently used as input for Multi-INTACT. For comparison, we also perform single-molecular trait INTACT analysis for expression and protein data, respectively.

We first assess the ability of the Multi-INTACT method to identify PCGs. Specifically, we evaluate the averaged power and false discovery rate at the target FDR control level of 5% across all simulated datasets for all methods. The results show that Multi-INTACT exhibits optimal power while properly controlling type I errors (Figure 2). Multi-INTACT outperforms existing methods that use a single gene product for PCG implication, including TWAS, colocalization analysis, and INTACT. We find that TWAS methods suffer from severely inflated type I errors due to failure to account for LD hitchhiking, while colocalization methods properly control type I errors but are overly conservative. Finally, while the single-trait INTACT method maintains fairly consistent power and FDR across TWAS prediction models (Table S2), Multi-INTACT achieves higher power mainly because additional gene products are considered.

**Figure 2:**
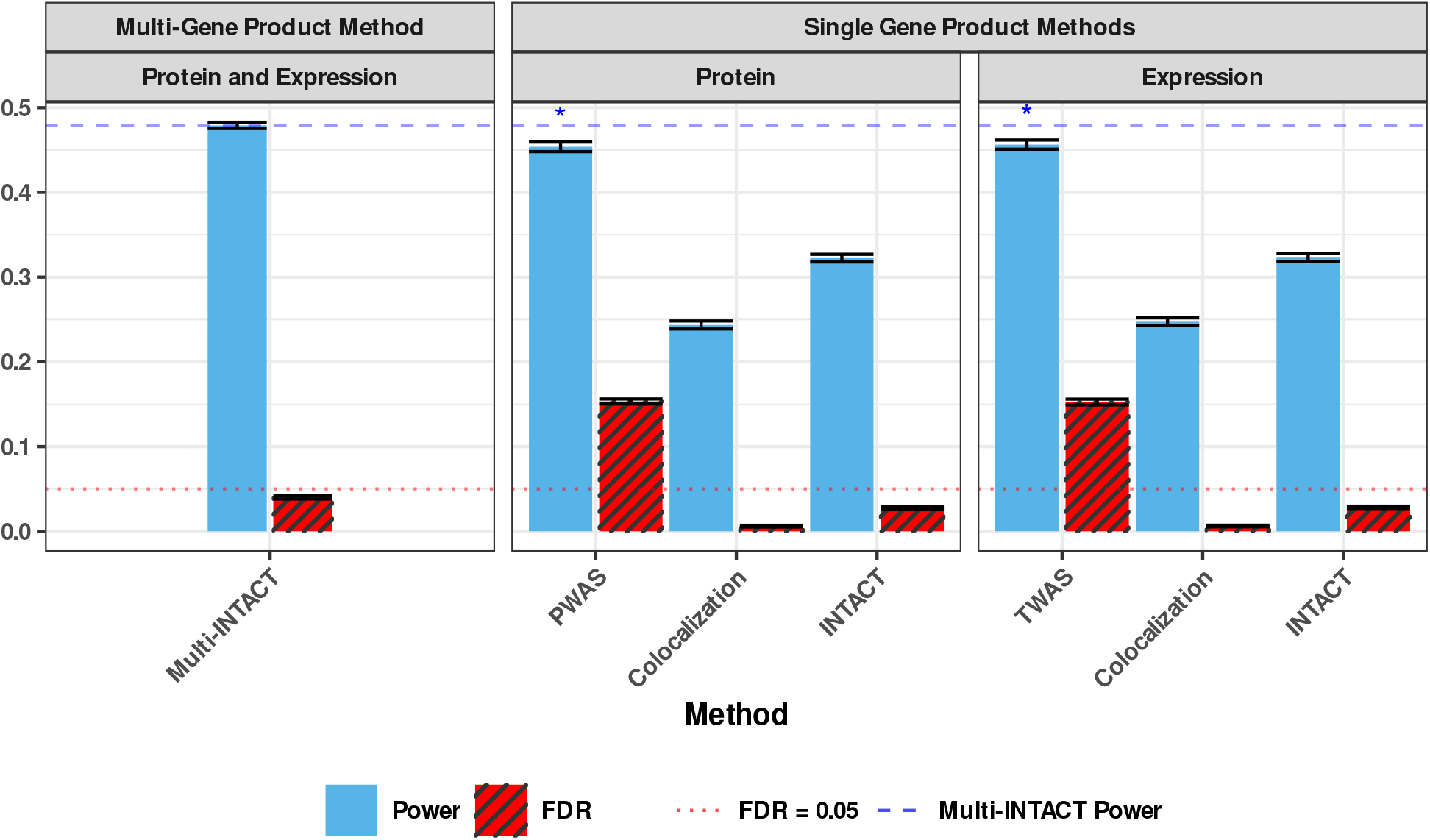
Average realized power and false discovery rates for each integrative PCG-implicating method at the 5% level over 100 simulated data sets. Methods are grouped by whether they consider multiple molecular phenotypes. Methods that consider only one molecular phenotype are grouped by omics platform. For ease of comparison, we include a dashed blue line to denote the Multi-INTACT power. Power columns representing methods with excessive false discoveries are marked with an asterisk. Error bars represent the standard error of the mean.

To illustrate Multi-INTACT’s ability to accommodate results from different TWAS prediction methods, we run Multi-INTACT using various popular molecular phenotype prediction models and observe that Multi-INTACT’s power and FDR results remain fairly consistent across different TWAS models (Table S3).

Additionally, we compute the gene probabilities of putative causality using only the summary association statistics of the simulated complex trait. We find the corresponding results are nearly identical to those obtained from individual-level GWAS data (Table S4). It should be acknowledged that our summary-level statistics represent a best-case scenario, as the LD information perfectly matches the underlying GWAS samples. In practice, LD information derived from a population reference panel often leads to imperfect characterization of sample LD and less-accurate inference results than what are obtained in this experiment.

Next, we evaluate Multi-INTACT as a means to identify relevant gene products for PCGs. To this end, we first apply the EM algorithm to estimate the proportions of PCG mechanisms in each simulated dataset. Then, we compute the corresponding gene-level posterior model probabilities for *M_E_, M_P_,* and *M_E_*_+_*_P_* . We find that the EM algorithm estimates of the mechanism distributions are reasonably accurate, i.e., the true proportion of PCGs following a mechanism always falls within the interquartile range of the EM algorithm estimate distribution (Figure S2). The distributions of the posterior model probabilities, stratified by the underlying true causal mechanisms, are shown in Figures 3 and S3-S5. Although the data do not completely distinguish the true mechanism in all settings, we find that the true mechanisms are always assessed with the highest posterior probabilities on average. Finally, we compute the gene product relevance probabilities for *E* and *P* across all genes. Using the gene product relevance probabilities as classification scores, we generate cumulative true discovery versus false discovery curves (Figure 4). Multi-INTACT outperforms alternative methods that rely on a single molecular trait at a time in identifying relevant causal gene products from highly ranked gene candidates.

**Figure 3:**
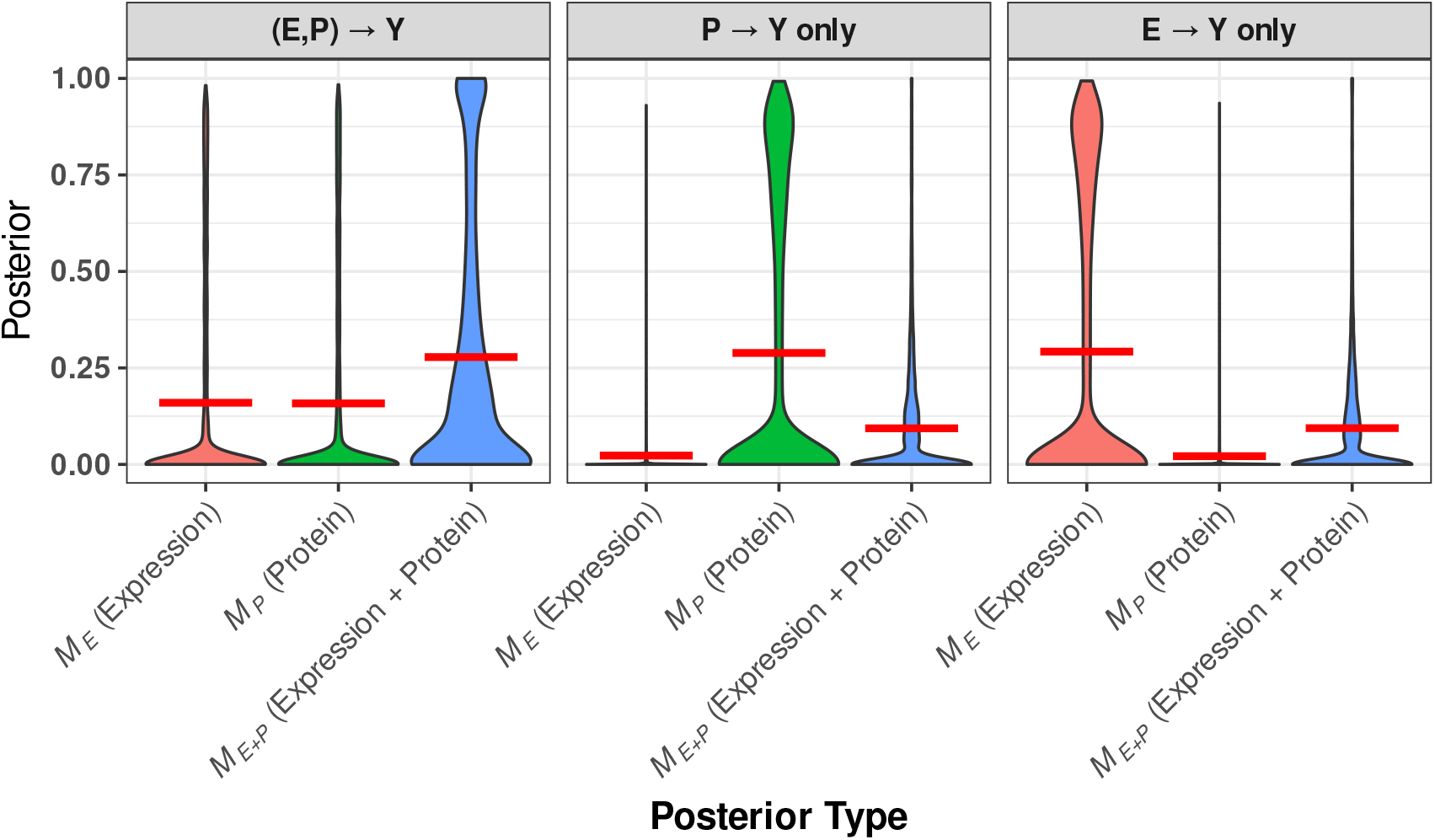
Distributions of posteriors for each gene product-to-trait effect scenario and three posterior types. The distributions represent genes that have nonzero causal effects on Y (through at least one of expression or protein). For each violin plot, a horizontal red line denotes the mean of the distribution.

**Figure 4:**
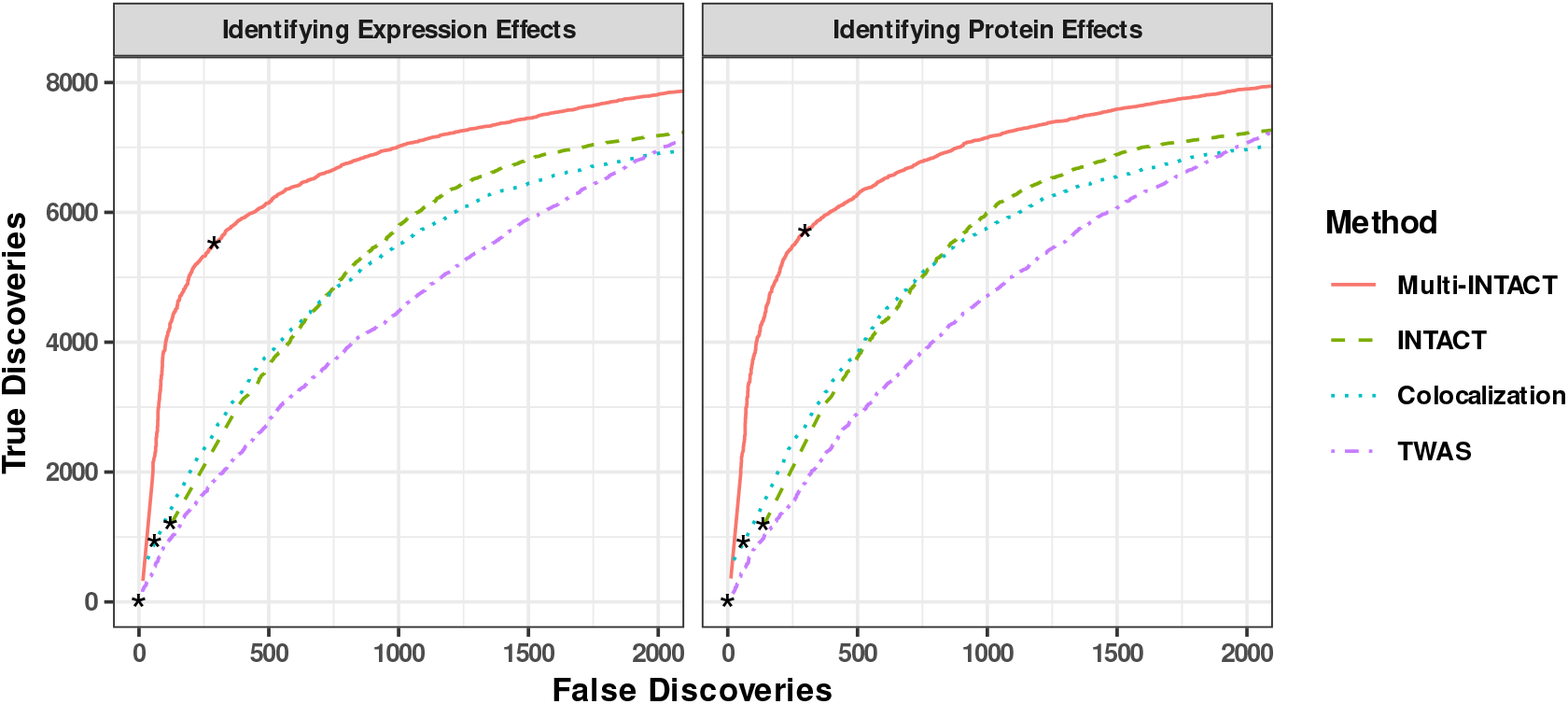
True discovery versus false discovery curves comparing the performances of Multi-INTACT and INTACT for classifying gene-to-trait effects. Curves are truncated at a false discovery rate threshold of 0.20 for Multi-INTACT in order to represent the highest-ranked genes. Results represent 100 simulated data sets with causal genes from a variety of possible DAGs. For Multi-INTACT, we use gene product relevance probabilities *P* (*E |* data) and *P* (*P |* data) as scores for detecting *E → Y* and *P → Y* effects, respectively. We use the INTACT posterior, fastENLOC gene-level colocalization probability, and PTWAS z-score magnitude with the respective molecular phenotype (expression, left; protein, right) as a classification score in each panel for comparison. For each curve the false discovery rate threshold of 0.05 is denoted by an asterisk.

### 2.3 Analysis of METSIM Metabolon Metabolite GWAS Data

To demonstrate the advantages of jointly considering multiple gene product molecular phenotypes, we apply INTACT and the proposed Multi-INTACT procedures to 1408 Metabolic Syndrome in Men (METSIM) study plasma metabolite GWASs [45], integrating the UK Biobank pQTL data [4] and multi-tissue GTEx eQTL data (v.8) [3] to identify PCGs and putative biological mechanisms. Details of the analysis, including algorithms used for TWAS and colocalization, are discussed in Methods.

The METSIM Metabolon metabolite study includes 10,188 Finnish men from Kuopio examined from 2005 to 2010 [45]. Study participants are whole-genome sequenced, yielding *>*26 million represented variants that pass QC procedures. For a description of the METSIM data pre-processing, refer to Supplemental Methods.

Our previous work highlights the discordance, or the lack of inferential reproducibility, in implicating PCGs when colocalization and TWAS analyses are applied to gene expression data [8, 20]. To determine whether a similar pattern holds in applications of proteomics data, we first perform UK Biobank pQTL-metabolite GWAS integrative analysis and compare the implicated PCGs (at 5% FDR level) from proteome-wide association study (PWAS), colocalization, and INTACT analyses across the 1408 metabolites. PWAS identifies 2,217 genes, colocalization analysis identifies 170 genes, and INTACT, combining colocalization and PWAS evidence, identifies 293 PCGs. Our results imply that *∼*95% of the PWAS genes do not show colocalization evidence, suggesting that most of these findings are likely due to the LD hitchhiking effects previously discussed in the context of TWAS analysis. Although INTACT is effective at guarding against LD hitchhiking, its statistical power is compromised and can be improved by incorporating more relevant gene products.

Next, we apply Multi-INTACT, integrating the UK Biobank pQTL data and the eQTL data representing one (at a time) of 49 tissues from the GTEx project. We first compute the gene probabilities of putative causality and identify PCGs at 5% FDR level separately in each metabolite-tissue pair. The full Multi-INTACT results from this analysis are summarized in the supplemental data. Among the tested 68,992 tissue-metabolite pairs, Multi-INTACT identifies 8,610 PCG-tissue-metabolite triplets, notably more than the expression-only or protein-only INTACT analyses which implicate 4,128 and 5,682 triplets, respectively. Upon stratifying the results by tissues, it is clear that although the number of discoveries varies by tissue, Multi-INTACT consistently implicates more genes than the expression-only and protein-only INTACT analyses (Figure 5). The increase in the discoveries illustrates the improved power of combining relevant gene products. Additionally, while a large proportion of the triplets identified by Multi-INTACT are also implicated by at least one of the INTACT analyses, there are many PCGs identified only by Multi-INTACT (Figure S6).

**Figure 5:**
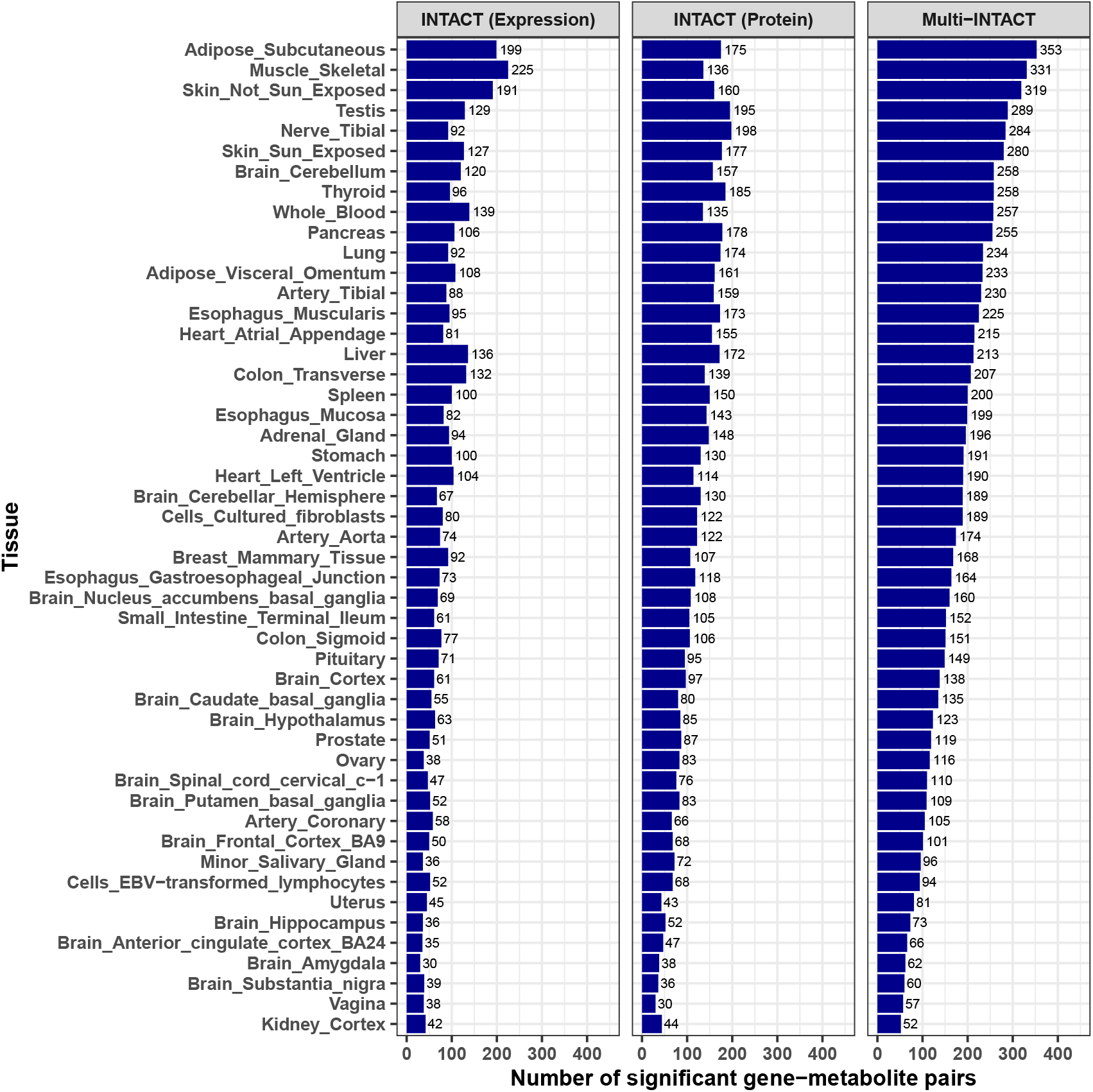
Multi-INTACT PCG implication results summary, by tissue. For each tissue-specific analysis, only genes with both expression and protein data are tested.

The overlap of PCG-metabolite pairs identified between tissues varies widely across the 49 GTEx tissues (Figure 6a). Unsurprisingly, Multi-INTACT results derived from brain cerebellum and cerebellar hemisphere expression data share a high proportion of identified PCG-metabolite pairs. Of the 894 unique PCG-metabolite pairs discovered in at least one tissue, we find that 23% of the PCG-metabolite pairs are implicated in a single tissue, and 50% of pairs are implicated in between 2 and 10 tissues (Figure 6b). Meanwhile, 4% of PCG-metabolite pairs are identified in *>*40 tissues. We find that the discrepancy of PCG discovery across tissues is driven by the tissue-dependent variability of eQTL discovery, which has been observed in tissue-specific TWAS analysis and attributed to both variation in statistical power and to biological factors, such as tissue-specific gene regulation. As a result, many genes do not have expression prediction models in all tissues. Here, we caution against interpreting tissue-specific PCG discoveries solely by biological (or statistical) factors.

**Figure 6:**
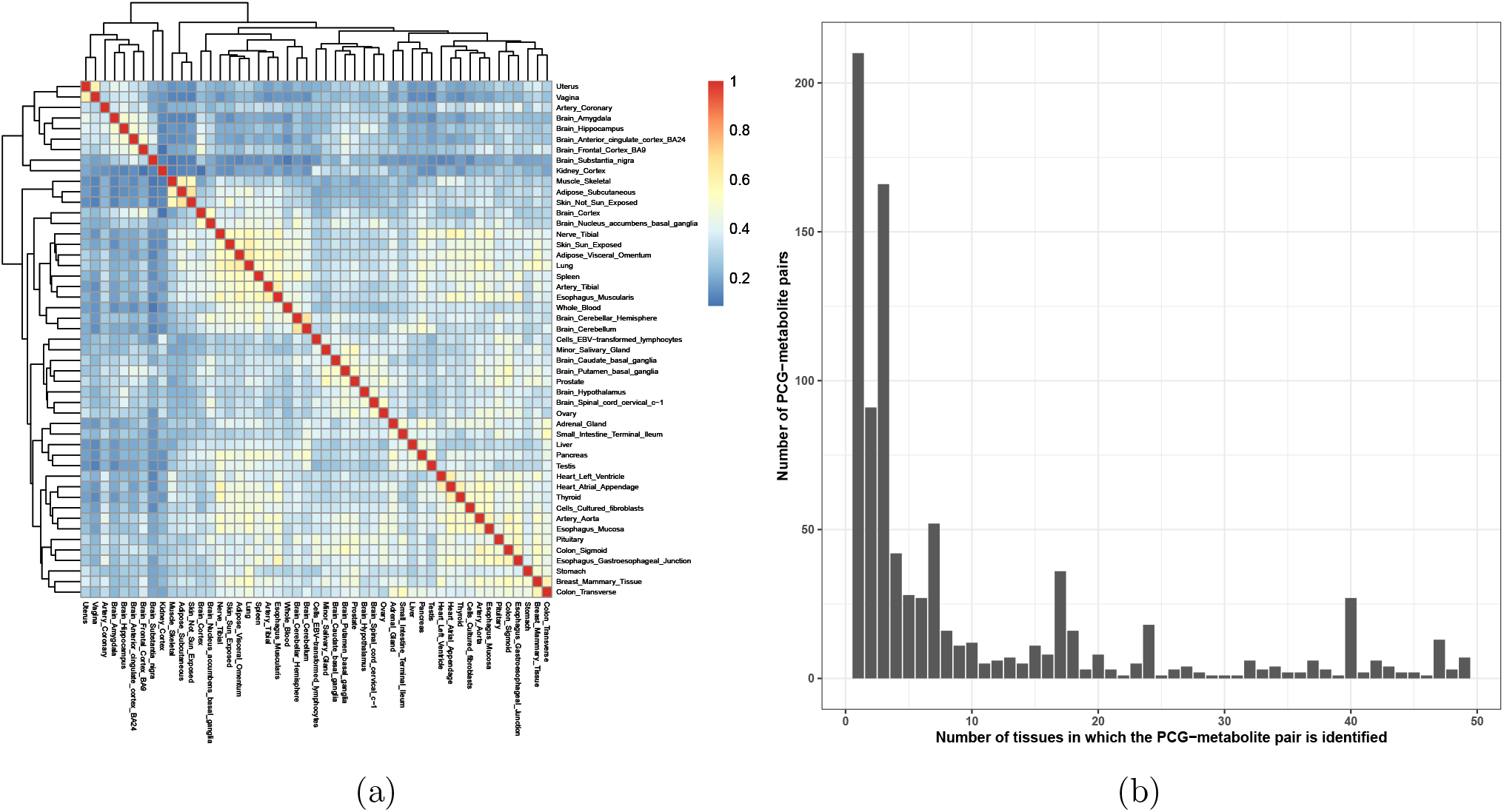
Overlap between gene-metabolite pairs discovered by Multi-INTACT using expression data across GTEx tissues. (a), Pairwise overlap of PCG-metabolite pairs implicated across tissues. Fill color denotes (# gene-metabolite pairs implicated by both tissues)/(# gene-metabolite pairs implicated by at least one of the tissues). (b), The distribution of the number of tissues in which Multi-INTACT identifies gene-metabolite pairs.

To validate the Multi-INTACT PCG findings, we compare our inferences to a high-quality annotated causal gene set for a group of metabolites by a knowledge-based approach (KBA). The KBA nominates PCGs by matching known metabolite biochemistry to functions of genes near strong GWAS signals [38, 39, 46, 47]. For our validation analysis, we limit the KBA nominations to genes that have both pQTLs and eQTLs in at least one tissue (i.e., candidates for Multi-INTACT analysis), where the KBA nominates 423 unique gene-metabolite pairs in total. The overlapping of the PCGs implicated by each integrative approach and the KBA are summarized in Figures 7. The expression-only and protein-only analyses identify 238 and 184 known gene-metabolite KBA pairs, respectively. In contrast, Multi-INTACT results overlap with 304, or *∼*70%, of the annotated KBA pairs, demonstrating superior power over the existing methods.

**Figure 7:**
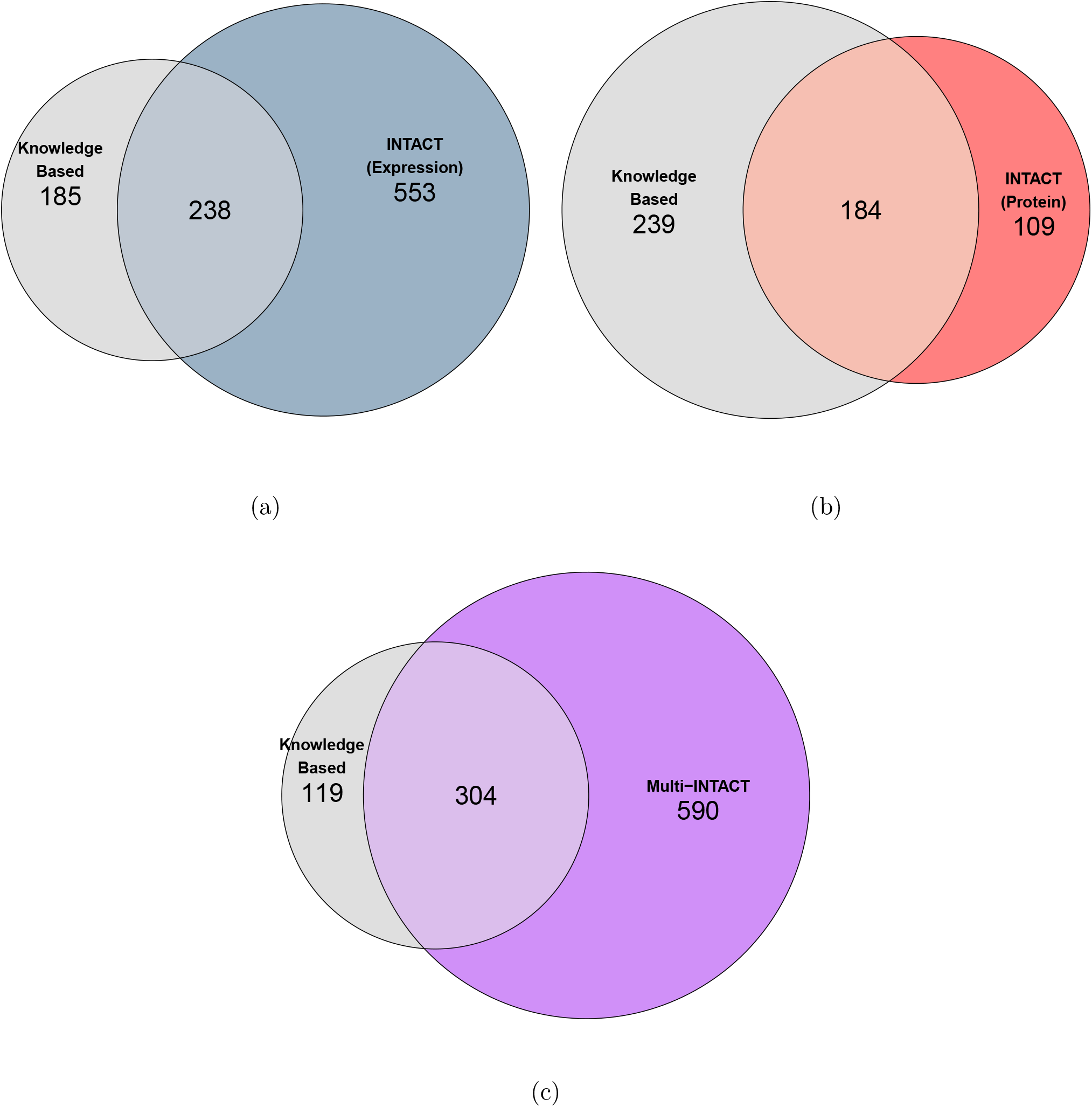
Overlap between gene-metabolite pairs discovered by integrative approaches and the knowledge-based approach (KBA) (a), INTACT using GTEx multi-tissue eQTL data. Numbers represent pairs implicated in at least one tissue. (b), INTACT using the UK Biobank pQTL data set. (c), Multi-INTACT using both GTEx and UK Biobank QTL data sets. There are 289 unique pairs implicated by KBA and either INTACT-expression or INTACT protein. Of these pairs, 287 are represented in the intersection of the KBA and Multi-INTACT discoveries.

Although the KBA is intended to benchmark Multi-INTACT, it does not provide an exhaustive list of all biologically-feasible PCG-metabolite pairs. Some pairs implicated by Multi-INTACT, but not the KBA, may reflect known biology. For example, we identify *GSTA1* as a PCG for DHEA-S (C100000792) in both liver and adrenal cortex tissues. This finding may reflect previous biological evidence that both *GSTA1* and DHEA-S have roles in metabolizing prostaglandins [48, 49].

Next, we investigate the directional consistency of different gene product-to-trait effects from the PCGs implicated by Multi-INTACT. The central dogma states that the flow of genetic information from DNA to RNA to protein is one-directional, implying that the sign of a genes effect should be the same across gene products. Despite this, many studies [50, 51, 52, 53] report complex relationships between expression levels, protein abundance, and disease risks. Although Multi-INTACT is not specifically designed to estimate gene product-to-trait effects, the signed TWAS and PWAS test statistics from *confidently inferred PCGs* should represent their qualitative directional effects. PCGs are confidently inferred if they are statistically significant based on gene probability of putative causality at 5% FDR level. In this analysis, we further select a subset of Multi-INTACT gene-tissue-metabolite triplets for which both gene expression and protein abundance are deemed relevant gene products. To this end, we focus on the set of triplets whose gene product relevance probabilities for both expression and protein are *≥* 0.50. For this selected set of triplets, we examine the directional consistency of the *z*-statistics from the corresponding TWAS and PWAS analyses.

Overall, of the 8,007 analyzed triplets, 5,000 (*∼* 62%) show matching directions from both gene products, while 3,007 (*∼* 38%) show opposite directions. Interestingly, some tissues known to play key roles in metabolism show high directional consistency. For example, the highest directional-consistency proportion is observed in liver (Figure 8), where 88% expression-to-trait and protein-to-trait effects are concordant and significantly higher than the remaining tissue “population” (p-value = 5.2 *×* 10*^−^*^12^). The concordant proportion increases to 96% when intersecting the Multi-INTACT results with the KBA results. Finally, we compute the Spearman correlation coefficient [54] to compare the signed log p-values of the TWAS and PWAS analyses. For the triplets implicated by Multi-INTACT, the correlation estimate (0.319, p-value = 2.18 × 10*^−^*^188^) is higher than the “population” average for tested metabolite-tissue pairs (mean = 0.156, variance = 0.002). A recent study comparing PWAS and TWAS in for blood lipid traits reports a similar range of correlation estimates as our population average [55].

**Figure 8:**
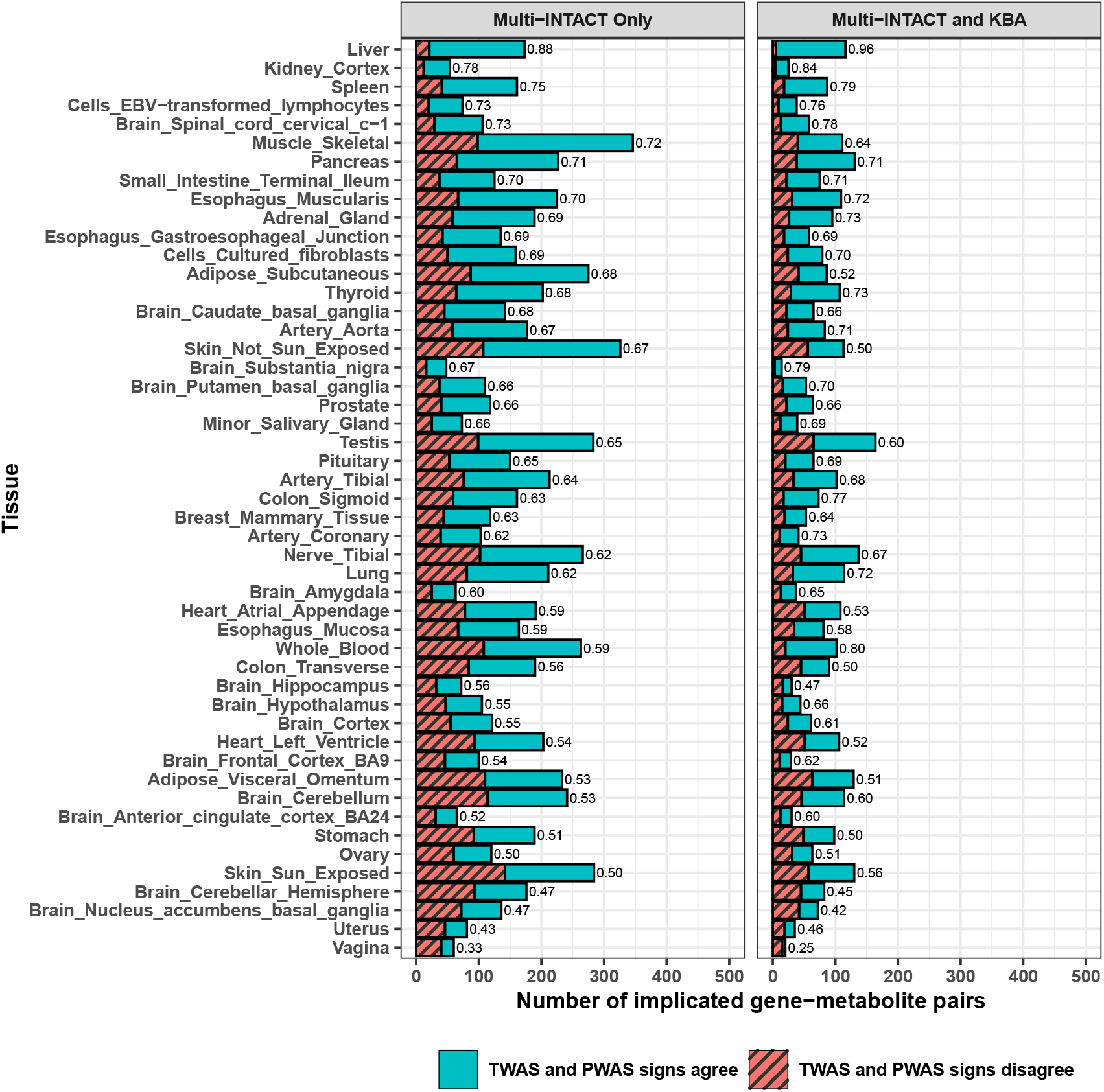
Gene-metabolite pairs with Multi-INTACT gene product relevance probabilities greater than 0.5 for both expression and protein, by tissue, with proportions of PWAS and TWAS test statistics that were the same/opposite signs. In the right panel, we show the intersection of genes implicated by Multi-INTACT and the knowledge-based approach (KBA) results. For each tissue-specific analysis, only genes with both expression and protein data are tested.

Lastly, we demonstrate the utility of Multi-INTACT for gene set enrichment analysis (GSEA). We focus our analysis on Multi-INTACT results derived from lipid metabolite data and liver expression data based on known biology [56] and results from previous analyses. For each target gene tested among the liver-lipid metabolite results, we form an aggregated probability of putative causality by combining the gene probability of putative causality across metabolites (see Methods for details). We then apply INTACT-GSE [8], a recently introduced GSEA method. We use the aggregated probability of putative causality as input for INTACT-GSE. These probabilities have the unique advantage of quantifying the uncertainty of the presence/absence of a PCG for at least one lipid metabolite, which is a key to obtaining unbiased enrichment estimates for candidate gene sets. We examine 213 Biological Process (BP) GO terms, estimating enrichment and the corresponding 95% confidence intervals. In summary, we identify 16 GO terms that are nominally significant at the 5% level, i.e., their confidence intervals do not overlap with 0 (Table 1). Among these findings, cholesterol metabolic process (GO:0008203) steroid metabolic process (GO:0008202) chemical reactions and pathways involving fatty acids (GO:0006631) are strongly enriched and reflect the well-known roles of liver in human metabolism. Full INTACT-GSE results are available in the supplemental data.

**Table 1:** Top enriched Biological Process GO terms from the INTACT-GSE gene set enrichment analysis. Each listed term is nominally significant at the 5% level.

| Biological Process<br>GO Term ID | Term | INTACT-GSE<br>Enrichment Estimate<br>(log odds ratio) | 95% Confidence<br>Interval |
| --- | --- | --- | --- |
| GO:0008203 | cholesterol metabolic process | 3.069 | (1.907, 4.231) |
| GO:0008202 | steroid metabolic process | 2.633 | (1.609, 3.657) |
| GO:0006644 | phospholipid metabolic process | 2.141 | (1.057, 3.225) |
| GO:0006629 | lipid metabolic process | 2.026 | (1.045, 3.007) |
| GO:0042632 | cholesterol homeostasis | 2.018 | (0.840, 3.197) |
| GO:0006631 | fatty acid metabolic process | 1.997 | (0.931, 3.064) |
| GO:0006082 | organic acid metabolic process | 1.978 | (1.025, 2.930) |
| GO:0016042 | lipid catabolic process | 1.913 | (0.854, 2.972) |
| GO:0055088 | lipid homeostasis | 1.894 | (0.737, 3.051) |
| GO:0046395 | carboxylic acid catabolic process | 1.893 | (0.842, 2.943) |
| GO:0009636 | response to toxic substance | 1.852 | (0.715, 2.989) |
| GO:0006805 | xenobiotic metabolic process | 1.805 | (0.520, 3.091) |
| GO:0001889 | liver development | 1.616 | (0.333, 2.899) |
| GO:0007041 | lysosomal transport | 1.616 | (0.333, 2.899) |
| GO:0007034 | vacuolar transport | 1.485 | (0.223, 2.747) |
| GO:0042594 | response to starvation | 1.373 | (0.124, 2.621) |

## 3 Discussion

We present Multi-INTACT, a novel statistical method for mechanism-aware PCG implication by integrating GWAS and multiple types of molecular QTL data. Through simulations and real data analysis, we demonstrate that Multi-INTACT properly controls type I errors by effectively guarding against LD hitchhiking and identifies causal genes with increased statistical power compared to other popular PCG implication methods. Additionally, its ability to simultaneously evaluate the relevance of multiple gene products (i.e., those exerting direct effects on complex traits) can help gain insights on detailed molecular mechanisms of complex diseases, serving as a potential tool for drug target discovery.

We motivate and derive the Multi-INTACT method under a non-parametric graphical model representing the established IV analysis with multiple endogenous variables, where the model assumptions focus on the conditional independence relationships rather than the distributional assumptions on molecular and complex phenotypes. Consequently, Multi-INTACT is robust and capable of handling various types of (e.g., quantitative, binary, and categorical) phenotypic data. Furthermore, we show that with additional linearity and normality assumptions, the Multi-INTACT model can be represented by a structural equation model (Equation 2 in Methods), which naturally extends the SEMs previously used in integrative genetic association analysis with a single molecular phenotype [6, 5].

The unique inference strategy implemented in Multi-INTACT is to leverage colocalization evidence and prevent TWAS/PWAS association signals driven *solely* by pleiotropic effects. This is a key difference from the alternative strategy that attempts to explicitly estimate and control potential pleiotropic effects [5, 6]. In our numerical experiments and real data analysis, we find that the Multi-INTACT strategy is conceptually simple, computationally efficient, and practically effective. Nevertheless, we acknowledge that, despite many unresolved statistical and computational challenges, accurate estimation of pleiotropic effects has the potential to further increase the power of PCG implication. We will to explore the possibility of combining these strategies in future work. Additionally, the use of a multiplicative combination of colocalization and multivariable regression evidence (via Bayes rule) in Multi-INTACT inference can also be justified by Dempster-Shafer (DS) theory [57] as described in the original INTACT paper. Specifically, the combination of the colocalization and regression evidence is an application of Dempster’s rule of combination. This application is a generalization of the INTACT evidence integration step in which each evidence source integrates information across multiple gene products rather than only the transcriptome.

In our joint analysis of METSIM Metabolon metabolite GWAS, GTEx eQTL, and UK Biobank pQTL data, we highlight the practical utility of Multi-INTACT, showing its potential in validating known and uncovering unknown molecular mechanisms of complex traits. Our investigation of directional consistency of gene-to-trait effects of PCGs between different gene products illustrates the complexity of the underlying scientific problem: although the observed patterns are sensible and expected by existing theoretical and experimental evidence, it is considerably challenging to interpret these findings for individual gene-tissue pairs. The observational association data have intrinsic limitations for further explorations. We hope that our findings can serve as a starting point for careful design of experimental investigations. In addition, we demonstrate that Multi-INTACT output can be directly applied to gene set enrichment analysis, a key feature to further validate and explore PCG findings. With improved information from multiple gene products, gene set enrichment analysis becomes more powerful compared to our previous analysis using a single gene product [8].

Although Multi-INTACT offers multiple improvements over existing PCG implication techniques, it does have limitations. The input from the existing colocalization and TWAS/PWAS analysis methods and the quality of currently available genetic data ultimately impact the performance of the Multi-INTACT analysis. Future advances in methods development and data generation in the related areas will help improve Multi-INTACT analysis. Other limitations that Multi-INTACT shares with many integrative analysis methods include a reliance on prior biological knowledge to determine relevant tissues or cell types and a focus on model selection/testing rather than estimating gene-to-trait effects. Finally, Multi-INTACT explicitly focuses on gene products. Although this feature makes implicating PCGs more direct and interpretable, there is room for improvement by incorporating some indirect but potentially important genomic information (e.g., chromatin structure, methylation status, and 3D genome structure). Under the Bayesian framework of Multi-INTACT, it seems feasible to consider additional genomic information by modifying the prior formulation. We will address these challenges in our future work.

## 4 Conclusion

In conclusion, Multi-INTACT integrates multiple gene products to reliably infer causal genes and mechanisms. Importantly, Multi-INTACT assesses which molecular gene products are the most relevant to disease, providing insights into possible drug targets. The Multi-INTACT software implementation is computationally efficient and can be applied using summary statistics of GWAS data. While we show that Multi-INTACT is useful for integrating expression and protein QTL datasets, our method has strong potential for studying complex trait etiology as the availability of molecular QTL datasets increases.

## 5 Methods

### 5.1 Structural Equation Model for Multi-INTACT

The Multi-INTACT method can be represented by a structural equation model (SEM), extending the SEM for TWAS analysis [8]. Consider a sample of *N* individuals from a single cohort. For a target gene, let ***E***(*N ×* 1), ***P*** (*N ×* 1), and ***G***(*N × p*) denote their expression levels, protein abundance, and genotypes for *p cis* genetic variants, respectively. The measurements of the complex trait of interest and the unobserved confounding are represented by ***Y*** (*N ×* 1) and ***U*** (*N ×* 1). All observed phenotype measurements are assumed to be pre-centered. Let *p*-vectors ***β****_E_* and ***β****_p_* denote the genetic effects on respective gene products for all *cis*-variants of the target gene and the *p*-vector ***β****_Y_* represents potential pleiotropic effects. Finally, we denote the *E → Y* and *P → Y* effects by *γ* and *δ*, which are of interest for inference. Reflecting the graphical model in Figure 1b, the proposed SEM is given by

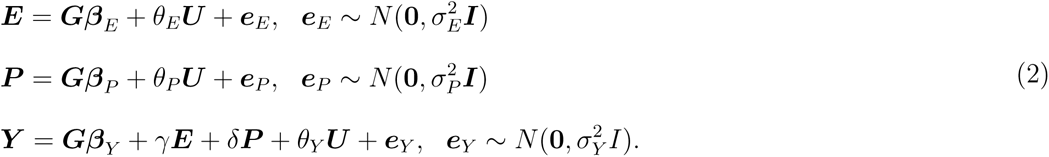

Noticeably, the above SEM does not explicitly specify potential *E → P* and/or *P → E* effects. Implicitly, these effects are accounted for by the corresponding residual error terms, i.e., ***e****_E_* and ***e****_P_* . The SEM is also similar to the multivariable Mendelian Randomization (MVMR) models discussed in genetic epidemiology [42, 58, 59, 60]. Their major difference lies in the inference strategy.

To assess a target gene for its putative causality, we consider testing a null hypothesis,

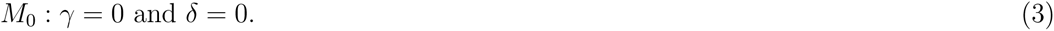

In Multi-INTACT, we adopt the Bayesian strategy of model selection and assess the posterior probability of *M*_0_. The strategy also naturally extends to the subsequent task of assessing relevant gene products by evaluating (and marginalizing from) the posterior probabilities of the alternative models,

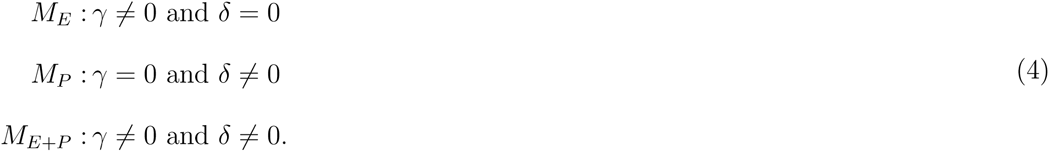

Note that the main goal of Multi-INTACT is to distinguish between different models representing different mechanisms for a candidate gene. These models are characterized by *M*_0_*, M_E_, M_P_*, and *M_E_*_+_*_P_* . Our goal is fundamentally different from rigorously estimating the causal effects *γ* and *δ*. This important point dictates our formulation of the statistical problems and the inference procedure for fitting SEM (2).

A key feature of the Multi-INTACT method is the use of colocalization evidence to guard against widespread LD hitchhiking. This feature is motivated by the following observation from the proposed SEM: if a gene product has a non-zero effect on the complex trait of interest (e.g., *δ /*= 0), its causal molecular QTL must also impose a non-zero genetic effect on the complex trait (e.g., *β_p_ · δ /*= 0). That is, colocalization is a necessary condition for *γ /*= 0 or *δ /*= 0 under the proposed model. In comparison, TWAS and PWAS signals driven by LD hitchhiking are *not* expected to exhibit colocalization evidence. However, in current practice, colocalization analysis is often severely underpowered [19]. Acknowledging this caveat, instead of requiring all implicated PCGs to show a high level of colocalization evidence, our default implementation of Multi-INTACT essentially filters out candidate genes that lack even modest colocalization evidence.

To compute the likelihood, we simplify SEM 2 for inference. Because the confounding ***U*** is unobserved, its effects on ***E***, ***P***, and ***Y*** are absorbed into the respective residual error terms. Consequently, ***E*** and ***P*** become endogenous variables (under the one-sample design) in the final regression equation of ***Y*** . To properly examine the gene-to-trait effects *γ* and *δ*, Multi-INTACT constructs two genetic instruments,

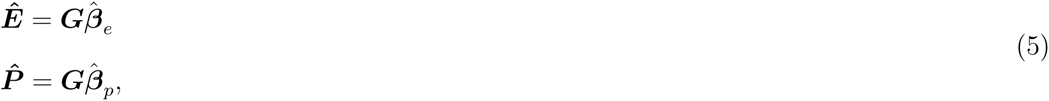

such that they are uncorrelated with ***U*** and ***e****_Y_* . Furthermore, as the colocalization prior effectively controls for the pleiotropic effects, we choose not to explicitly estimate ***β****_Y_* and absorb its effect into the residual error ***e****_Y_* . In the end, we fit the following regression model to compute the marginal likelihood for *γ /*= 0 or *δ /*= 0, i.e.,

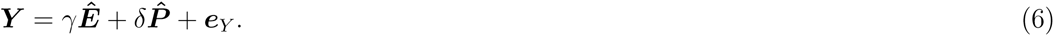

Note that, the estimated percentage of variance explained (PVE), or *R*^2^, by fitting (6) is identical to the squared canonical correlation between ***Y*** and (***Ê***, ***P̂***), which we reason from the perspective of multivariable IV analysis without the SEM formulation in Results section.

Because the regression model (6) requires only genetically predicted molecular gene products, the inference procedure can be naturally extended to multi-sample designs, in which the prediction models for ***Ê*** and ***P̂*** are learned in a cohort different from the GWAS samples. Importantly, the extension to data from multi-sample designs does not alter the causal implication of the original SEM model.

### 5.2 Computing Gene Probability of Putative Causality

We compute the gene probability of putative causality for a target gene by applying the Bayes rule,

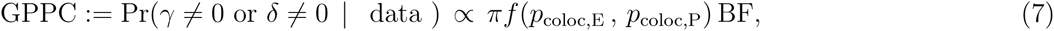

where *p*_coloc,E_ and *p*_coloc,P_ denote the pre-computed gene-level colocalization probabilities of respective molecular QTLs and GWAS hits, *πf* (*p*_coloc,E_, *p*_coloc,P_) denotes the composite prior probability of putative causality, and BF represents the marginal likelihood/Bayes factor. See Supplemental Methods for details on the computation of BF, estimation of *π*, and the prior function *f* .

### 5.3 Computing Gene Product Relevance Probability

The computation of gene product relevance probabilities breaks down to evaluating posterior probabilities for *M_E_, M_P_*, and *M_P_* _+_*_E_*. To be consistent with the GPPC calculation, we specify Pr(*M*_0_) = 1 *− πf* (*p*_coloc,E_, *p*_coloc,P_) and define the following conditional priors for the three alternative models,

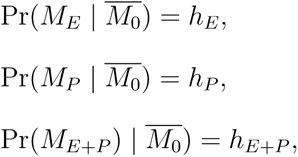

where 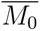 indicates the set of non-null models and *h_E_* + *h_P_* + *h_E_*_+_*_P_* = 1.

Following an empirical Bayes procedure, we first compute the Bayes factor of each non-null model for all target genes and design an EM algorithm to obtain the MLE of (*h_E_, h_P_, h_E_*_+_*_P_*) by pooling all candidate genes (Supplemental Methods). We then evaluate the required posterior model probabilities by plugging the estimated hyperparameters and subsequently compute the gene product relevance probabilities for each target gene using Equation (1).

### 5.4 Computation with GWAS Summary Statistics

The Multi-INTACT computation can be approximated using summary statistics of GWAS data. At a minimum, single-SNP association testing *z* scores, weights for molecular phenotype prediction, and an appropriate LD reference panel are required to compute multivariate Wald statistic for Bayes factor calculation. We show the details of two approximate computation methods using the minimum GWAS summary statistics in Supplemental Methods. The same information can also be used to compute gene-level colocalization evidence.

It is worth emphasizing that summary statistics-based computation is not exact, and the loss of accuracy is expected. In practice, we find that if the LD reference panel matches well with the underlying GWAS samples, the results approximate the exact computation (using individual-level data) well (Figure S7). Most importantly, there should not be inflation of type I errors in PCG implication due to replacing individual-level data with the corresponding summary statistics under this setting. However, the consequence of severe mismatch between the LD panel and GWAS samples is unclear and needs further investigation.

### 5.5 Simulation study

We use genotypes of 477K SNPs on chromosome 5 from 500 GTEx samples, including 1198 consecutive genes, each with at least 1500 common *cis*-SNPs. The complex and molecular phenotypes are simulated based on the complete DAG models shown in the second row of Table S1, all of which assume no direct effects between *E* and *P* (Note that Multi-INTACT inference does not assume or use this information).

Specifically, in each simulated data set, each of 1198 genes’ causal mechanisms is independently drawn from a Multinomial(*π*_0_,*π_E_*,*π_P_*,*π_E_*_+_*_P_*) distribution, where *π*_0_, *π_E_*, *π_P_*, and *π_E_*_+_*_P_* represent probabilities of the null, expression-only, protein-only, and expression-and-protein models. For each gene, we randomly select two eQTLs and two pQTLs, where one variant is both a causal eQTL and pQTL. We select a distinct causal GWAS SNP. All effect sizes (represented by edges in the DAGs) are drawn from a *N* (0*, φ*^2^) distribution, with *φ* set to 0.6 and residual error variances set to 1 to yield realistic signal-to-noise ratios. The distribution of the proportion of variance explained for all simulated phenotypes is shown in Figure S8. The Mean PVE for gene expression, protein levels, complex trait, expression-mediated complex trait, and protein-mediated complex trait are 0.167, 0.166, 0.159, 0.050, and 0.049, respectively. These values reasonably resemble the observed data in practice.

We simulate 100 datasets using the above scheme by varying the values of (*π*_0_,*π_E_*,*π_P_*,*π_E_*_+_*_P_*), with approximately 1/3 simulated datasets taking values from (0.8,0.1,0.05,0.05), (0.8,0.05,0.1,0.05), and (0.8,0.05,0.05,0.1), respectively. We use the different simulated datasets to examine the accuracy of the estimated (*h_E_*, *h_P_*, *h_E_*_+_*_P_*) values from the proposed EM algorithm. The FDR and power for PCG discovery are calculated across all simulated datasets.

In order to examine the robustness of Multi-INTACT in the presence of effects between expression and protein levels, we perform additional simulations to represent all 9 scenarios shown in Table S1. We describe the design of these additional simulations in Supplemental Methods. Power and FDR results for these simulations are shown in Figures S9 - S11.

### 5.6 Preprocessing of UK Biobank pQTL data and multi-tissue GTEx eQTL data

We use PTWAS [15] multi-tissue prediction models trained on the GTEx data set to predict expression for the individuals in the METSIM cohort. For PWAS analysis, we use the most significant cis-pQTL (+/- 1 Mb) to predict protein levels for each individual. We perform pair-wise colocalization analyses between the QTL data and metabolite GWASs using fastENLOC, computing gene level colocalization probabilities [20] to quantify the likelihood of a colocalized variant for each metabolite-transcript or metabolite-protein pair.

### 5.7 INTACT and Multi-INTACT analyses

We performed INTACT analyses using the default setting in the R package (linear prior and GLCP threshold *t* = 0.05). We focus on genes with both expression data and protein data available in the UK Biobank and GTEx prediction models. The number of tested genes per metabolite-tissue pair depends on the availability of expression and protein prediction models. The number of genes tested ranges from 186 (kidney cortex) to 1,795 (nerve tibial). The complete breakdown is shown in Figure S12.

### 5.8 INTACT-GSE pathway enrichment analysis for lipid metabolites

We perform probabilistic GSEA using the Multi-INTACT results derived from liver expression data. We use Multi-INTACT’s default settings (truncation threshold equal to 0.05 with the linear prior function) to compute posterior probabilities for each gene-metabolite triplet. For each tested gene, we compute an aggregated probability of putative causality across all lipid metabolites by

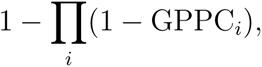

where GPPC*_i_* is the gene probability of putative causality for the examined gene and the *i*th lipid metabolite. Intuitively, the aggregated quantity represents the probability of the target gene is a PCG for at least one metabolite. We examine all GO BP terms for which at least one annotated gene has a nonzero aggregated probability of putative causality. GSEA input, including gene-level aggregated probabilities of putative causality, is included in the supplemental data. We access the GO term annotation data via the R package org.Hs.eg.db (v3.17.0).

## 6 Declarations

### 6.1 Ethics approval and consent to participate

Not applicable

### 6.2 Consent for publication

Not applicable

### 6.3 Availability of data and materials

The datasets generated and/or analyzed during the current study are available at https://tinyurl.com/ye2aysvf. All analyses in this paper can be reproduced using the code at https://github.com/jokamoto97/multi_intact_paper. The Multi-INTACT software is accessible in the INTACT R package at https://github.com/jokamoto97/INTACT.

### 6.6 Authors’ contributions

JO, FL, RPG, HKI, JM, CB, EBF, ML, MB, and XW and conceived and designed the research. All authors performed the research. JO, XY, BR, JC, and XW analyzed the resulting data. JO and XW wrote the paper. All authors read and approved the final manuscript.

## Acknowledgements

Not applicable

## References

[1] Hindorff, L. A. et al. Potential etiologic and functional implications of genome-wide association loci for human diseases and traits. Proceedings of the National Academy of Sciences 106, 9362–9367 (2009).

[2] Visscher, P. M. et al. 10 years of gwas discovery: biology, function, and translation. The American Journal of Human Genetics 101, 5–22 (2017).

[3] Consortium, G. The gtex consortium atlas of genetic regulatory effects across human tissues. Science 369, 1318–1330 (2020).

[4] Sun, B. B. et al. Plasma proteomic associations with genetics and health in the uk biobank. Nature 1–10 (2023).

[5] Mancuso, N. et al. Probabilistic fine-mapping of transcriptome-wide association studies. Nature genetics 51, 675–682 (2019).

[6] Barfield, R. et al. Transcriptome-wide association studies accounting for colocalization using egger regression. Genetic epidemiology 42, 418–433 (2018).

[7] Yuan, Z. et al. Testing and controlling for horizontal pleiotropy with probabilistic mendelian randomization in transcriptome-wide association studies. Nature communications 11, 1–14 (2020).

[8] Okamoto, J. et al. Probabilistic integration of transcriptome-wide association studies and colocalization analysis identifies key molecular pathways of complex traits. The American Journal of Human Genetics 110, 44–57 (2023).

[9] Zhao, S., et al. Adjusting for genetic confounders in transcriptome-wide association studies leads to reliable detection of causal genes. bioRxiv (2022).

[10] Gamazon, E. R. et al. A gene-based association method for mapping traits using reference transcriptome data. Nature genetics 47, 1091–1098 (2015).

[11] Zhu, Z. et al. Integration of summary data from gwas and eqtl studies predicts complex trait gene targets. Nature genetics 48, 481–487 (2016).

[12] Gusev, A. et al. Integrative approaches for large-scale transcriptome-wide association studies. Nature genetics 48, 245–252 (2016).

[13] Giambartolomei, C. et al. Bayesian test for colocalisation between pairs of genetic association studies using summary statistics. PLoS genetics 10, e1004383 (2014).

[14] Pividori, M. et al. Phenomexcan: Mapping the genome to the phenome through the transcriptome. Science Advances 6, eaba2083 (2020).

[15] Zhang, Y. et al. Ptwas: investigating tissue-relevant causal molecular mechanisms of complex traits using probabilistic twas analysis. Genome biology 21, 1–26 (2020).

[16] Giambartolomei, C. et al. A bayesian framework for multiple trait colocalization from summary association statistics. Bioinformatics 34, 2538–2545 (2018).

[17] Hormozdiari, F. et al. Colocalization of gwas and eqtl signals detects target genes. The American Journal of Human Genetics 99, 1245–1260 (2016).

[18] Zhu, A. et al. Mrlocus: Identifying causal genes mediating a trait through bayesian estimation of allelic heterogeneity. PLoS genetics 17, e1009455 (2021).

[19] Hukku, A. et al. Probabilistic colocalization of genetic variants from complex and molecular traits: promise and limitations. The American Journal of Human Genetics 108, 25–35 (2021).

[20] Hukku, A., Sampson, M. G., Luca, F., Pique-Regi, R. & Wen, X. Analyzing and reconciling colocalization and transcriptome-wide association studies from the perspective of inferential reproducibility. The American Journal of Human Genetics 109, 825–837 (2022).

[21] Nica, A. C. et al. Candidate causal regulatory effects by integration of expression qtls with complex trait genetic associations. PLoS genetics 6, e1000895 (2010).

[22] Nicolae, D. L. et al. Trait-associated snps are more likely to be eqtls: annotation to enhance discovery from gwas. PLoS genetics 6, e1000888 (2010).

[23] He, B., Shi, J., Wang, X., Jiang, H. & Zhu, H.-J. Genome-wide pqtl analysis of protein expression regulatory networks in the human liver. BMC biology 18, 1–16 (2020).

[24] Ferkingstad, E. et al. Large-scale integration of the plasma proteome with genetics and disease. Nature Genetics 53, 1712–1721 (2021).

[25] Yao, D. W., Oconnor, L. J., Price, A. L. & Gusev, A. Quantifying genetic effects on disease mediated by assayed gene expression levels. Nature genetics 52, 626–633 (2020).

[26] Raj, T. et al. Integrative transcriptome analyses of the aging brain implicate altered splicing in alzheimers disease susceptibility. Nature genetics 50, 1584–1592 (2018).

[27] Sun, Q. et al. From gwas variant to function: A study of 148,000 variants for blood cell traits. Human Genetics and Genomics Advances 3, 100063 (2022).

[28] Gandal, M. J. et al. Transcriptome-wide isoform-level dysregulation in asd, schizophrenia, and bipolar disorder. Science 362, eaat8127 (2018).

[29] Kikuchi, R. et al. An antiangiogenic isoform of vegf-a contributes to impaired vascularization in peripheral artery disease. Nature medicine 20, 1464–1471 (2014).

[30] Scotti, M. M. & Swanson, M. S. Rna mis-splicing in disease. Nature Reviews Genetics 17, 19–32 (2016).

[31] van den Hoogenhof, M. M., Pinto, Y. M. & Creemers, E. E. Rna splicing: regulation and dysregulation in the heart. Circulation research 118, 454–468 (2016).

[32] Liu, E. Y., Cali, C. P. & Lee, E. B. Rna metabolism in neurodegenerative disease. Disease models & mechanisms 10, 509–518 (2017).

[33] Cooper, T. A., Wan, L. & Dreyfuss, G. Rna and disease. Cell 136, 777–793 (2009).

[34] Cheng, J. C., Moore, T. B. & Sakamoto, K. M. Rna interference and human disease. Molecular Genetics and Metabolism 80, 121–128 (2003).

[35] Cheng, Y., LeGall, T., Oldfield, C. J., Dunker, A. K. & Uversky, V. N. Abundance of intrinsic disorder in protein associated with cardiovascular disease. Biochemistry 45, 10448– 10460 (2006).

[36] Swindell, W. R. et al. Proteogenomic analysis of psoriasis reveals discordant and concordant changes in mrna and protein abundance. Genome medicine 7, 1–22 (2015).

[37] Sriwijitkamol, A. et al. Reduced skeletal muscle inhibitor of *κ*b*β* content is associated with insulin resistance in subjects with type 2 diabetes: reversal by exercise training. Diabetes 55, 760–767 (2006).

[38] Yin, X. et al. Genome-wide association studies of metabolites in finnish men identify disease-relevant loci. Nature communications 13, 1–14 (2022).

[39] Yin, X. et al. Integrating transcriptomics, metabolomics, and gwas helps reveal molecular mechanisms for metabolite levels and disease risk. The American Journal of Human Genetics (2022).

[40] Robertson, K. D. Dna methylation and human disease. Nature Reviews Genetics 6, 597–610 (2005).

[41] Wang, J. et al. Atac-seq analysis reveals a widespread decrease of chromatin accessibility in age-related macular degeneration. Nature communications 9, 1–13 (2018).

[42] Burgess, S. & Thompson, S. G. Multivariable mendelian randomization: the use of pleiotropic genetic variants to estimate causal effects. American journal of epidemiology 181, 251–260 (2015).

[43] VanderWeele, T. J., Tchetgen, E. J. T., Cornelis, M. & Kraft, P. Methodological challenges in mendelian randomization. *Epidemiology (Cambridge*, Mass.) 25, 427 (2014).

[44] Didelez, V. & Sheehan, N. Mendelian randomization as an instrumental variable approach to causal inference. Statistical Methods in Medical Research 16, 309330 (2007). URL 10.1177/0962280206077743.

[45] Laakso, M. et al. The metabolic syndrome in men study: a resource for studies of metabolic and cardiovascular diseases. Journal of lipid research 58, 481–493 (2017).

[46] Lotta, L. A. et al. A cross-platform approach identifies genetic regulators of human metabolism and health. Nature Genetics 53, 54–64 (2021).

[47] Stacey, D. et al. Progem: a framework for the prioritization of candidate causal genes at molecular quantitative trait loci. Nucleic acids research 47, e3–e3 (2019).

[48] Bogaards, J. J., Venekamp, J. C. & van Bladeren, P. J. Stereoselective conjugation of prostaglandin a2 and prostaglandin j2 with glutathione, catalyzed by the human glutathione s-transferases a1-1, a2-2, m1a-1a, and p1-1. Chemical research in toxicology 10, 310–317 (1997).

[49] Markiewicz, L. & Gurpide, E. C19 adrenal steroids enhance prostaglandin f2*α* output by human endometrium in vitro. American journal of obstetrics and gynecology 159, 500–504 (1988).

[50] Liu, Y., Beyer, A. & Aebersold, R. On the dependency of cellular protein levels on mrna abundance. Cell 165, 535–550 (2016).

[51] Vogel, C. & Marcotte, E. M. Insights into the regulation of protein abundance from proteomic and transcriptomic analyses. Nature reviews genetics 13, 227–232 (2012).

[52] Wu, L. et al. Variation and genetic control of protein abundance in humans. Nature 499, 79–82 (2013).

[53] Battle, A. et al. Impact of regulatory variation from rna to protein. Science 347, 664–667 (2015).

[54] Myers, L. & Sirois, M. J. Spearman correlation coefficients. Differences between. Encyclopedia of statistical sciences 12 (2006).

[55] Zhang, D. et al. Proteome-wide association studies for blood lipids and comparison with transcriptome-wide association studies. bioRxiv 2023–08 (2023).

[56] Nguyen, P. et al. Liver lipid metabolism. Journal of animal physiology and animal nutrition 92, 272–283 (2008).

[57] Dempster, A. P. Upper and Lower Probabilities Induced by a Multivalued Mapping. The Annals of Mathematical Statistics 38, 325 – 339 (1967). URL 10.1214/aoms/1177698950.

[58] Burgess, S., Dudbridge, F. & Thompson, S. G. Multivariable mendelian randomization: the use of pleiotropic genetic variants to estimate causal effects. American journal of epidemiology 181, 290–291 (2015).

[59] Sanderson, E., Davey Smith, G., Windmeijer, F. & Bowden, J. An examination of multivariable mendelian randomization in the single-sample and two-sample summary data settings. International journal of epidemiology 48, 713–727 (2019).

[60] Rees, J. M., Wood, A. M. & Burgess, S. Extending the mr-egger method for multivariable mendelian randomization to correct for both measured and unmeasured pleiotropy. Statistics in medicine 36, 4705–4718 (2017).

