## Supplemental Figures, Tables, and Methods for "Integrative analysis of the genome, transcriptome, and proteome identifies causal mechanisms of complex traits"

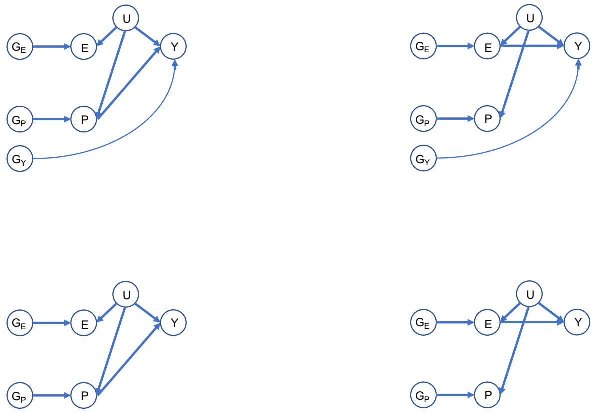


Figure S1: Various causal scenarios in which only one type of molecular QTL is colocalized with a causal GWAS hit. $G_E$, $G_P$, and $G_Y$ denote causal eQTLs, pQTLs, and distinct GWAS SNPs, respectively. $E$, $P$, and $Y$ denote gene expression, protein, and complex trait levels, respectively.

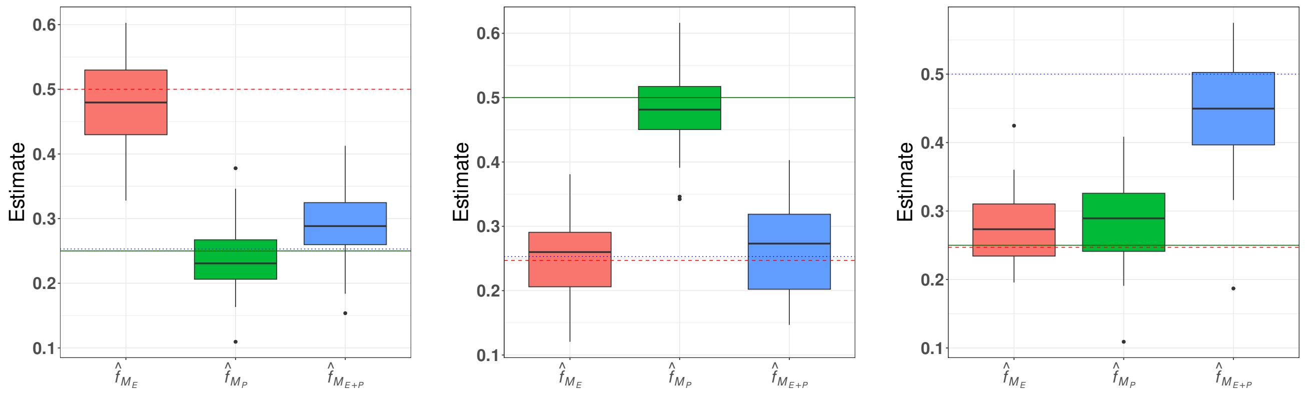


(a) (b) (c)

Figure S2: **EM algorithm estimate distributions across simulated data sets** Estimates are scaled to sum to 1. The dashed red line, solid green line, and dotted blue line denote the true proportion of $E \rightarrow Y$ causal genes, $P \rightarrow Y$ causal genes, and $(E, P) \rightarrow Y$ causal genes, respectively. Lines are slightly staggered in each plot for clarity.

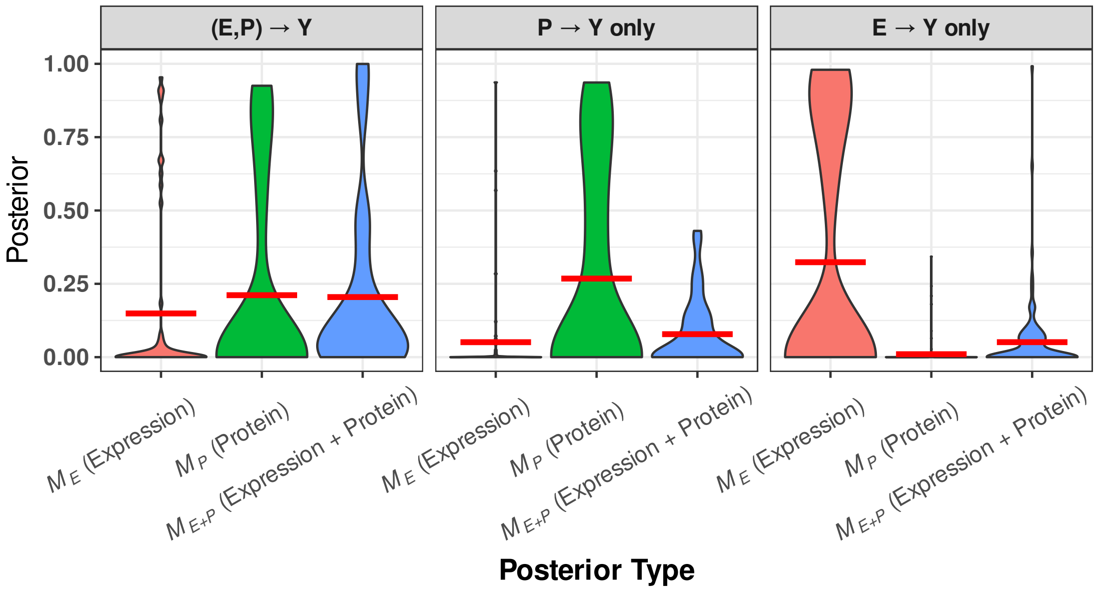


Figure S3: Distributions of posteriors for each gene product-to-trait effect scenario and three posterior types. The distributions represent genes that have nonzero causal effects on Y (through at least one of expression or protein). For each violin plot, a horizontal red line denotes the mean of the distribution. For this simulated data set, causal gene mechanisms were drawn from a $Multinomial(\pi_1, \pi_2, \pi_3) = Multinomial(0.5, 0.25, 0.25)$ distribution, where $\pi_1$ is the probability of $E \to Y$ *and* $P \to Y$ effects, $\pi_2$ is the probability of *only* $P \to Y$ effects, and $\pi_3$ is the probability of *only* $E \to Y$ effects.

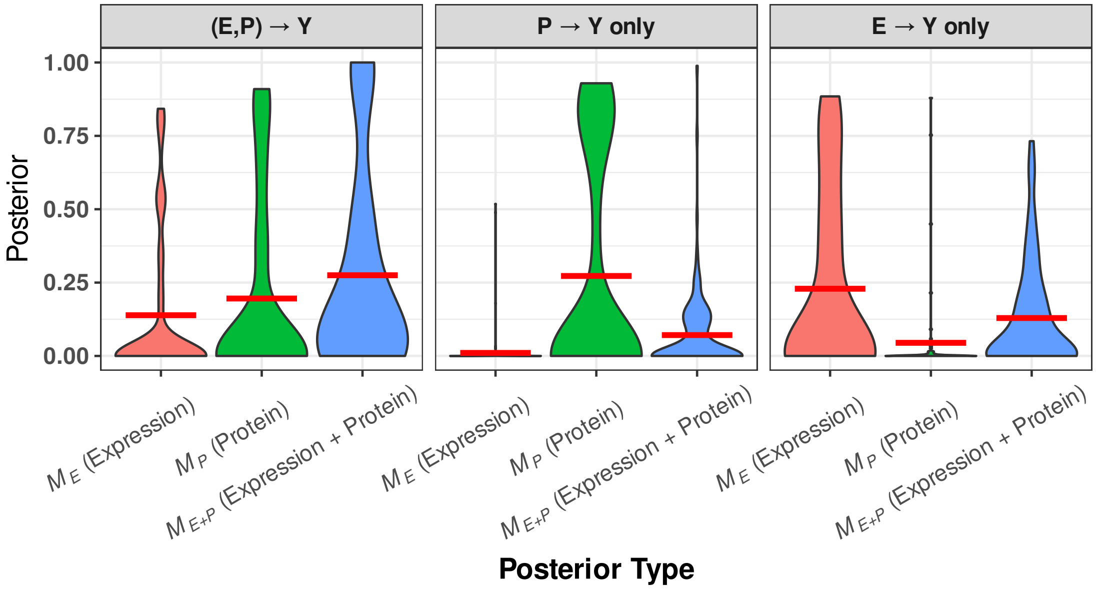


Figure S4: Distributions of posteriors for each gene product-to-trait effect scenario and three posterior types. The distributions represent genes that have nonzero causal effects on Y (through at least one of expression or protein). For each violin plot, a horizontal red line denotes the mean of the distribution. For this simulated data set, causal gene mechanisms were drawn from a $Multinomial(\pi_1, \pi_2, \pi_3) = Multinomial(0.25, 0.5, 0.25)$ distribution, where $\pi_1$ is the probability of $E \to Y$ *and* $P \to Y$ effects, $\pi_2$ is the probability of *only* $P \to Y$ effects, and $\pi_3$ is the probability of *only* $E \to Y$ effects.

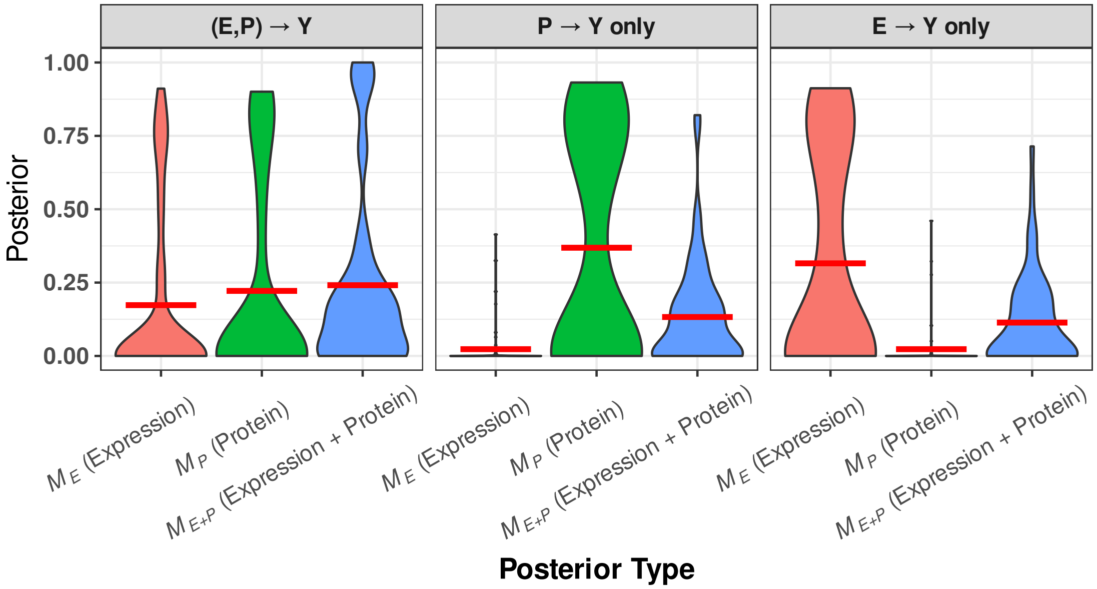


Figure S5: Distributions of posteriors for each gene product-to-trait effect scenario and three posterior types. The distributions represent genes that have nonzero causal effects on Y (through at least one of expression or protein). For each violin plot, a horizontal red line denotes the mean of the distribution. For this simulated data set, causal gene mechanisms were drawn from a $Multinomial(\pi_1, \pi_2, \pi_3) = Multinomial(0.25, 0.25, 0.5)$ distribution, where $\pi_1$ is the probability of $E \to Y$ *and* $P \to Y$ effects, $\pi_2$ is the probability of *only* $P \to Y$ effects, and $\pi_3$ is the probability of *only* $E \to Y$ effects.

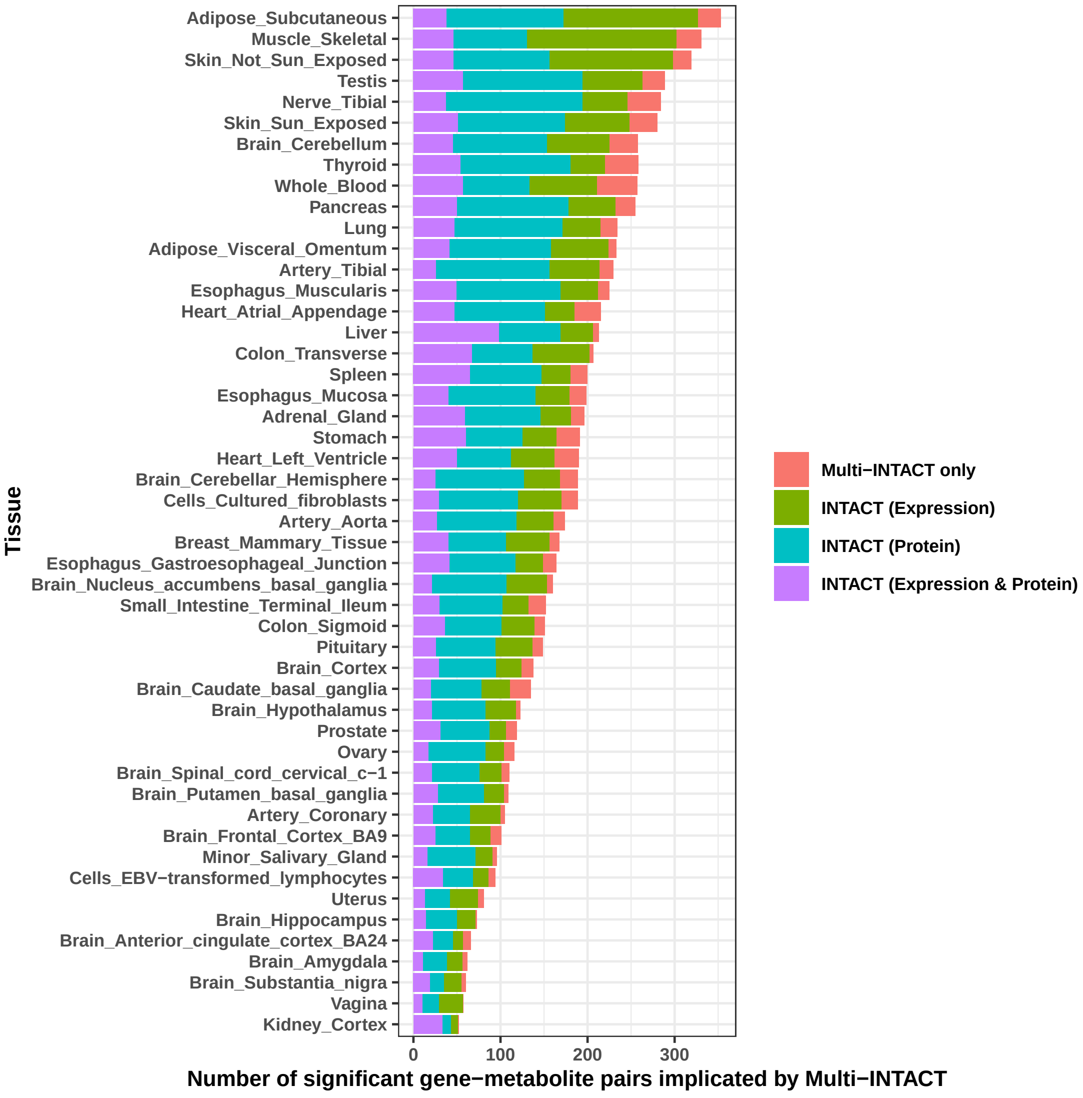


Figure S6: Multi-INTACT gene implication results, by tissue, with proportions implicated by marginal analyses. For each tissue-specific analysis, only genes with both expression and protein data are tested.

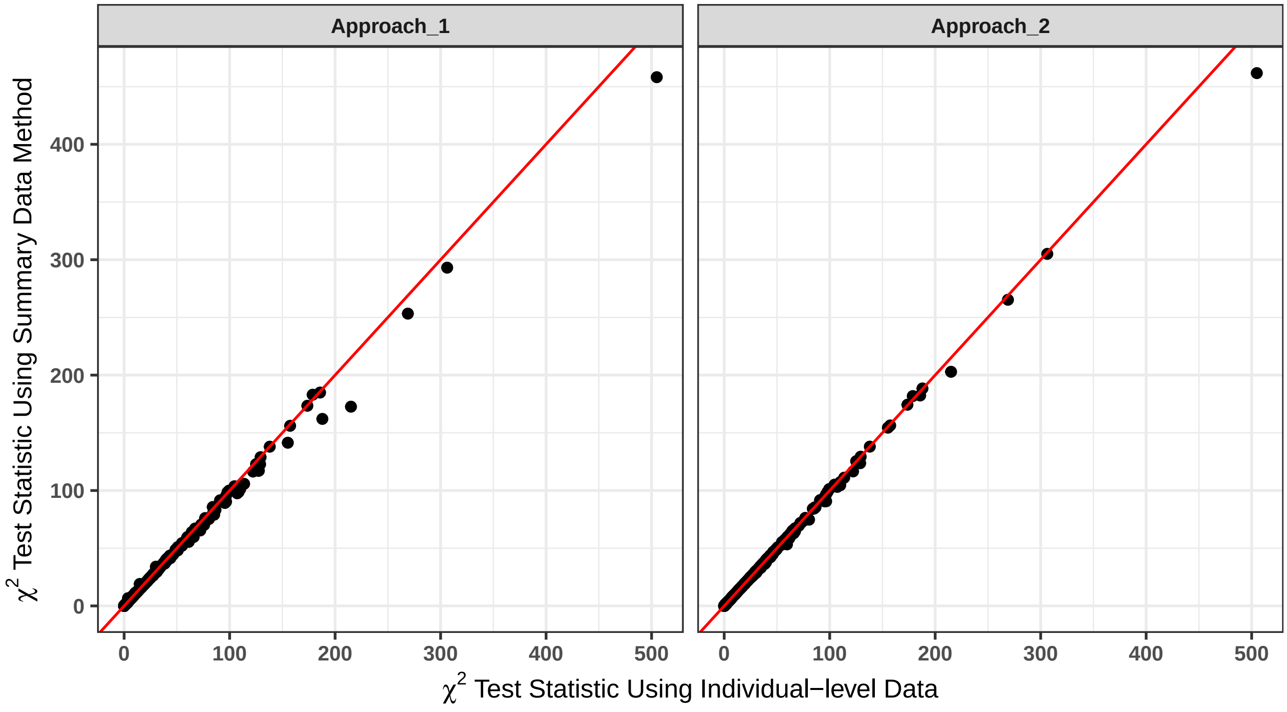


Figure S7: Comparison of two multivariate regression test statistic approximations based on summary level-data to the statistic computed from individual-level data. Approach 1 corresponds to the formula adapted from DAP-G, while Approach 2 corresponds to the MultiXcan formula. Descriptions of both approaches are provided in Supplemental Methods. A line at y = x is included in each panel for ease of comparison.

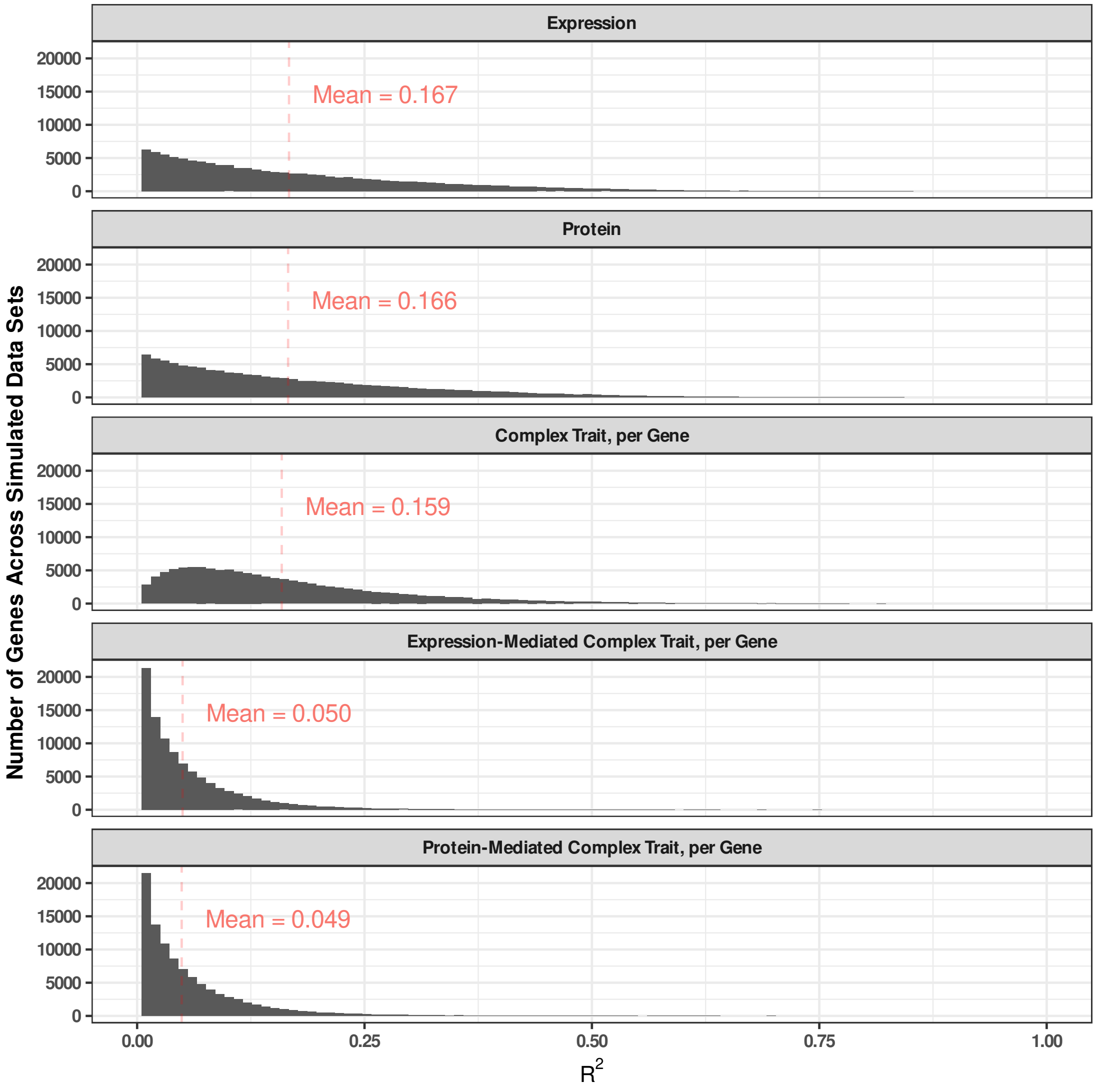


Figure S8: Distributions of expression heritability, protein heritability, complex trait heritability per gene, expression-mediated complex trait heritability per gene, and protein-mediated complex trait heritability per gene in 100 simulated data sets. Expression heritability is computed as the $R^2$ from the regression of expression on the true eQTLs $g_1$ and $g_2$. Similarly, protein heritability is the $R^2$ from the regression of protein on the true pQTLs $g_2$ and $g_3$. Complex trait heritability is computed as the $R^2$ from the regression of the complex trait on all QTLs in addition to the distinct GWAS SNP $g_4$. Expression-mediated complex trait heritability is computed as the $R^2$ from the regression of the complex trait on the causal eQTLs, while protein-mediated complex trait heritability uses the true pQTLs as covariates rather than eQTLs.

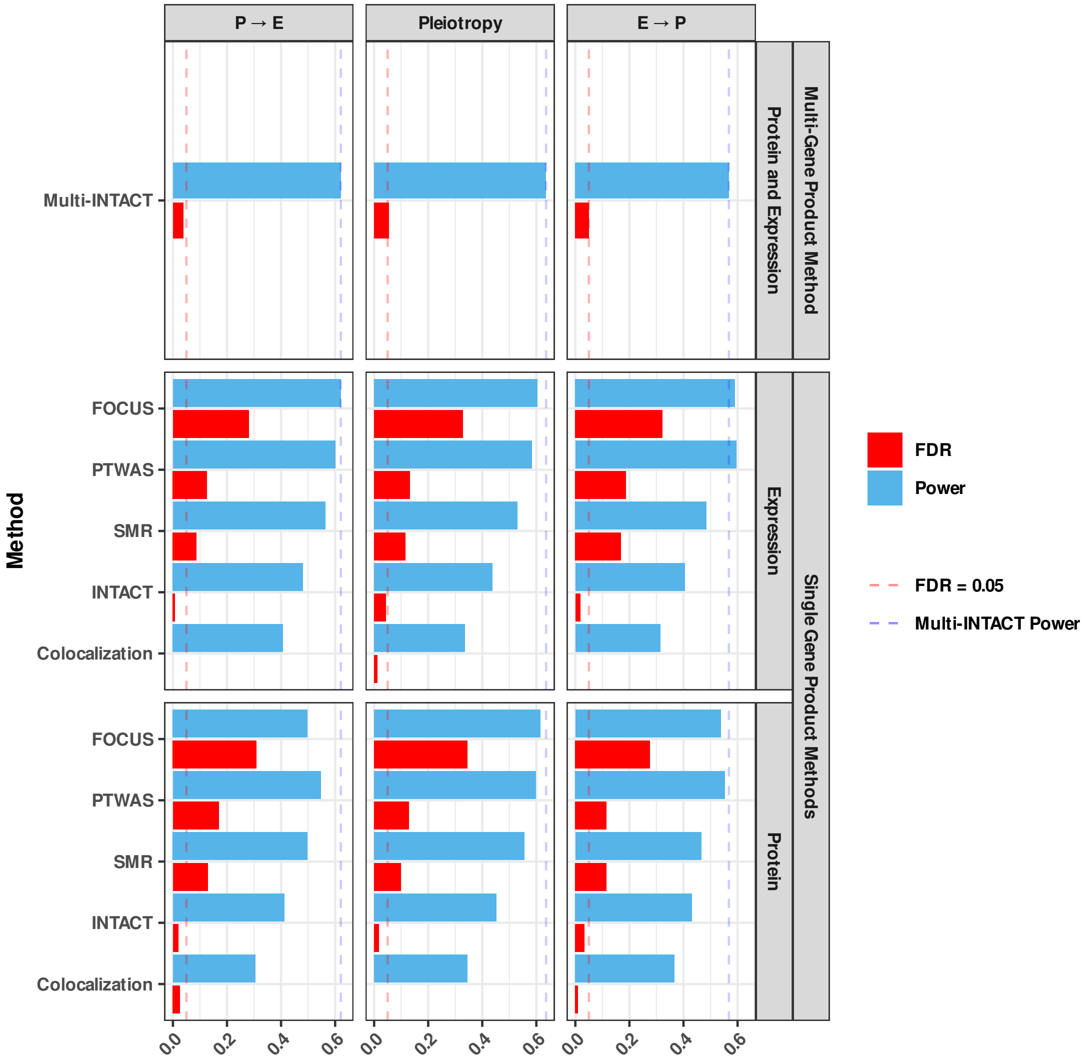


Figure S9: Multi-INTACT power and realized FDR for simulations in which the gene effects the complex trait though both protein and expression levels.

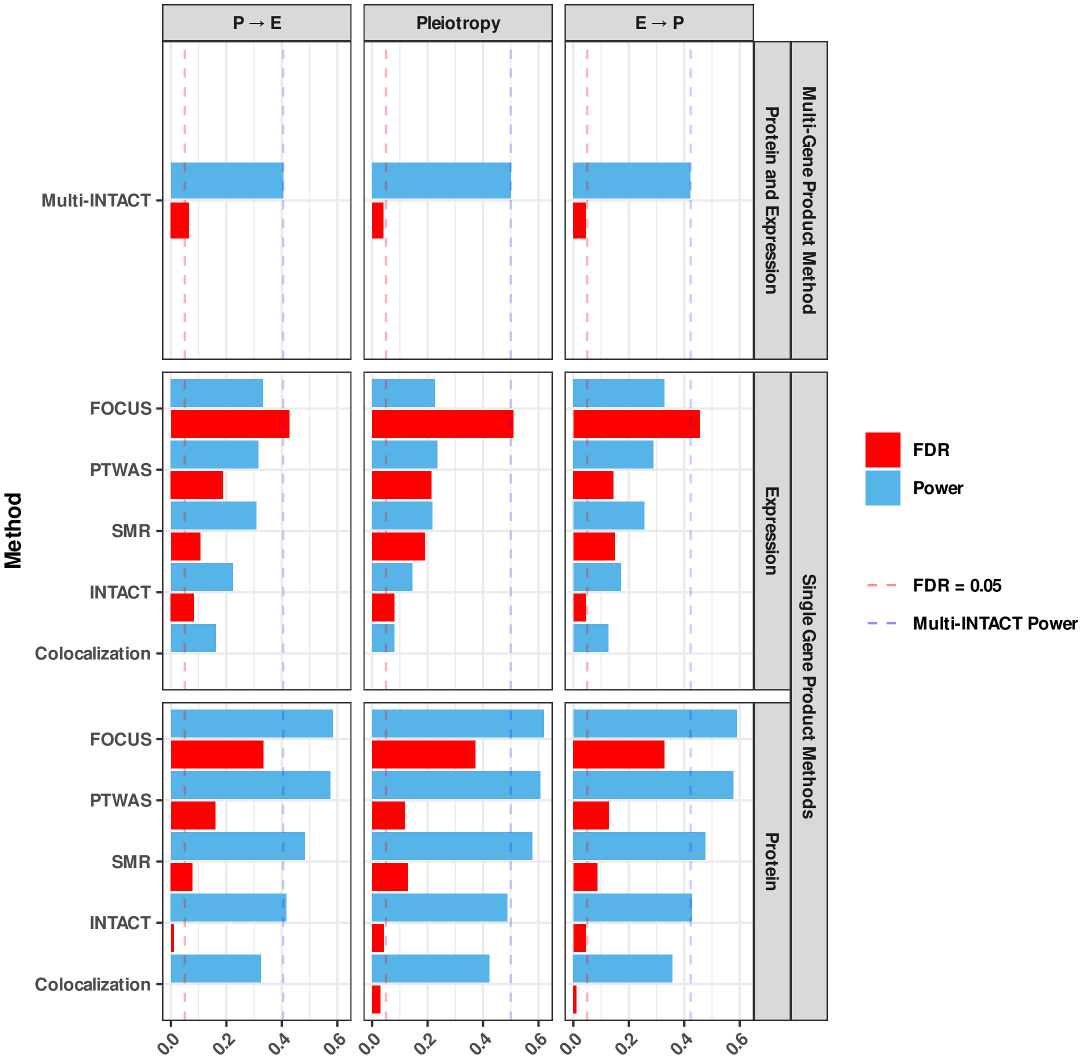


Figure S10: Multi-INTACT power and realized FDR for simulations in which the gene effects the complex trait though protein levels, but not expression levels.

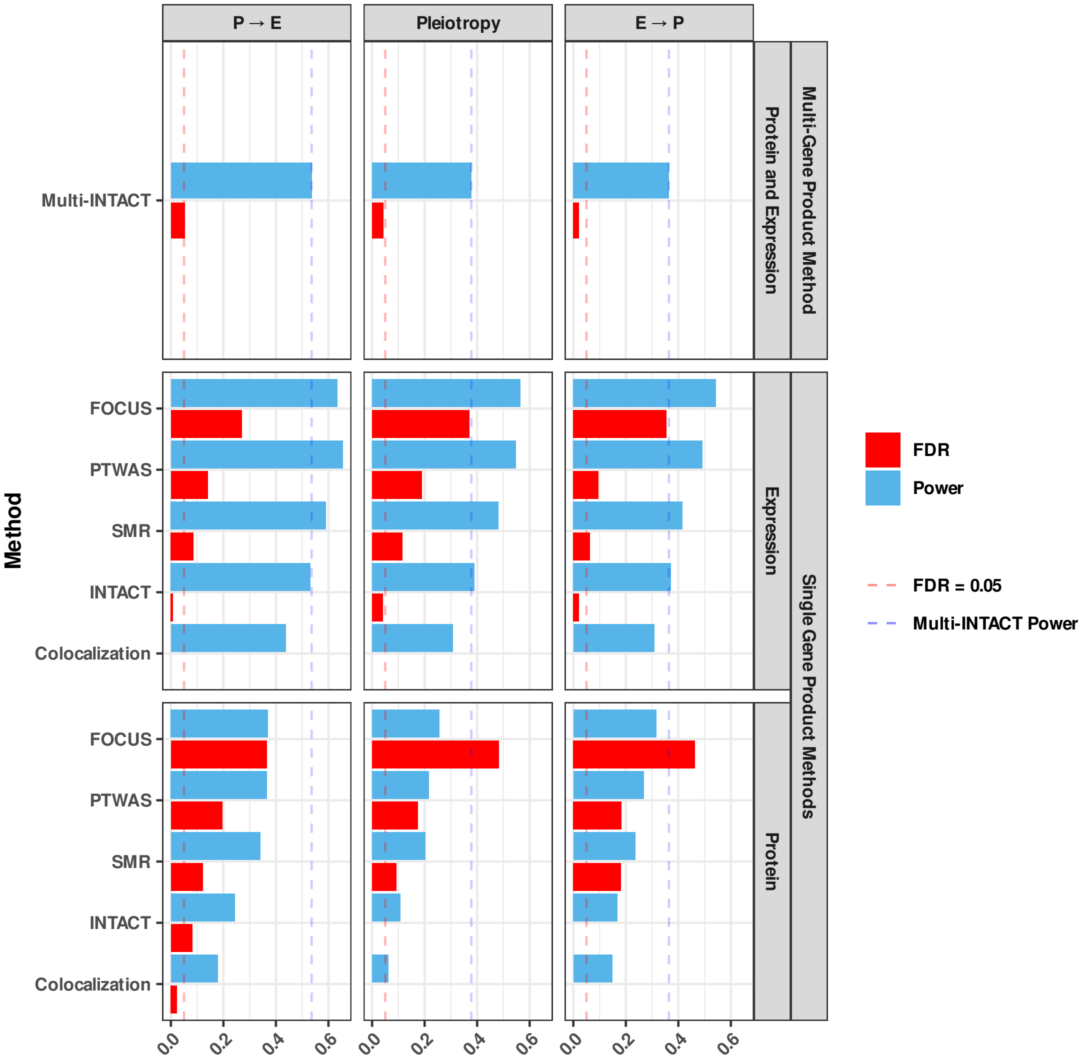


Figure S11: Multi-INTACT power and realized FDR for simulations in which the gene effects the complex trait though expression levels, but not protein levels.

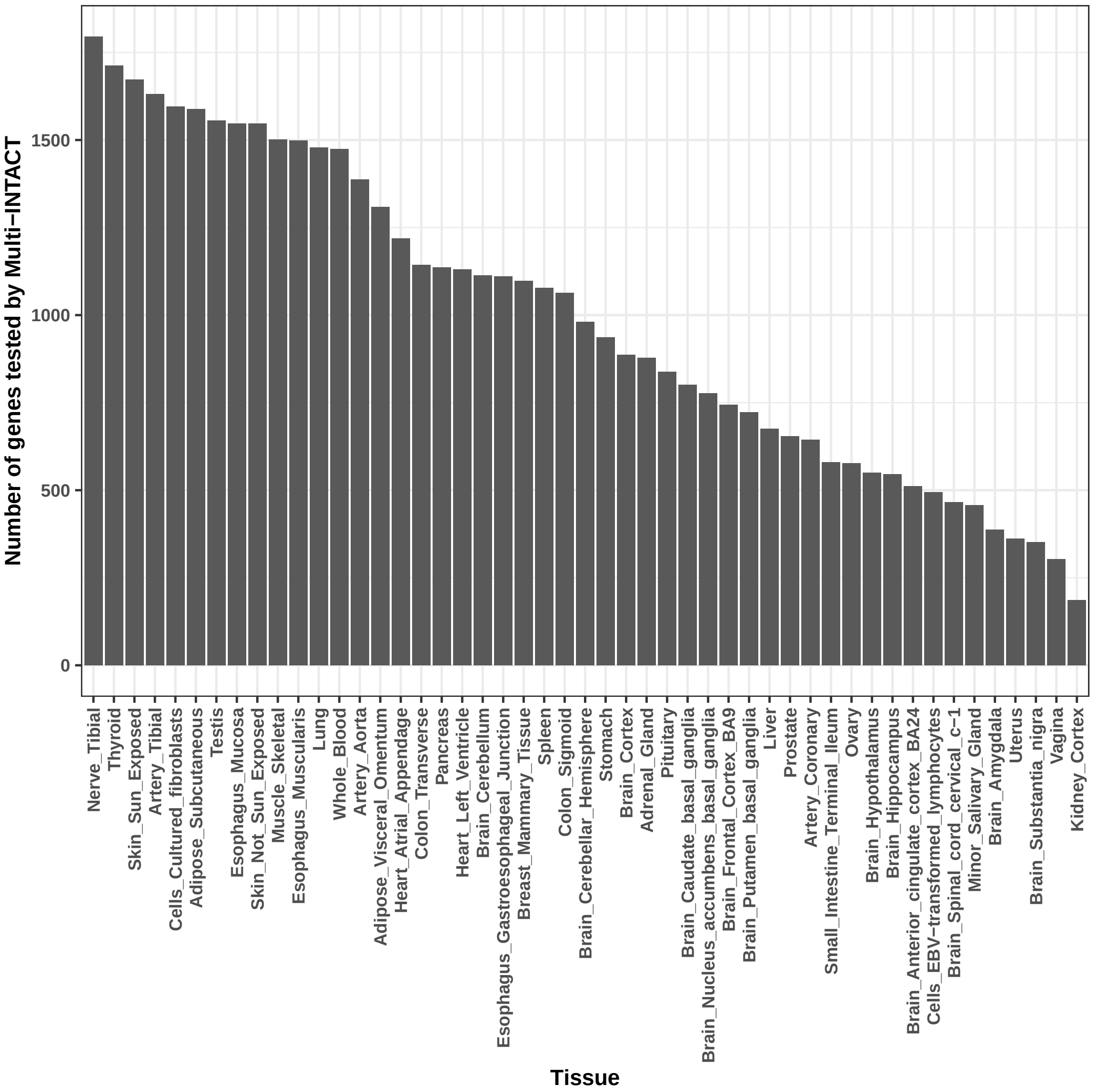


Figure S12: Number of genes tested by Multi-INTACT across tissues. Only genes with both expression and protein data are tested.

### Supplemental Tables

| | E→Y and P→Y effects | P→Y effect | E→Y effect |
|---|---|---|---|
| P→E effect | 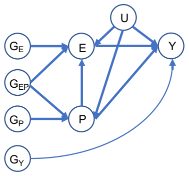 | 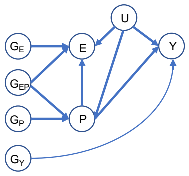 | 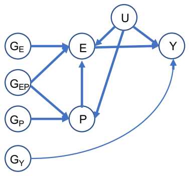 |
| No P→E or E→P effect | 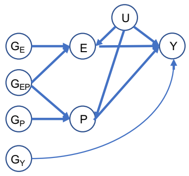 | 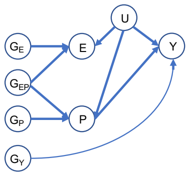 | 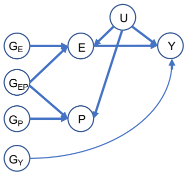 |
| E→P effect | 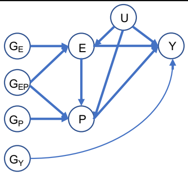 | 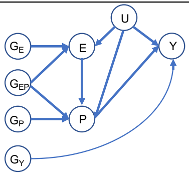 | 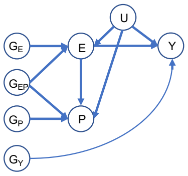 |

Table S1: Causal diagrams representing possible causal relationships between genotypes $G$, molecular traits $E$ and $P$, and a complex trait $Y$. Here, the genotypes are divided into those directly affect $P$ only ($G_P$), those directly affecting $E$ only ($G_E$), those directly affecting both $E$ and $P$ ($G_{EP}$), and those directly affecting $Y$ ($G_Y$).

| Molecular phenotype prediction model | Molecular phenotype | Power | FDR |
|---|---|---|---|
| PTWAS | Expression | 0.481 | 0.008 |
| | Protein | 0.412 | 0.020 |
| Single-variant | Expression | 0.469 | 0.000 |
| | Protein | 0.407 | 0.029 |
| Elastic net | Expression | 0.490 | 0.017 |
| | Protein | 0.416 | 0.019 |

Table S2: **Simulation INTACT results across a variety of expression and protein prediction models** For each simulated gene, TWAS and PWAS analyses are performed using the expression/protein prediction method indicated in the left-most column. The underlying causal DAG has $E \to Y$, $P \to Y$, and $P \to E$ effects. Realized power and false discovery rate (FDR) at the 5% control level are provided in the right-most columns.

| Molecular phenotype prediction model | Power | FDR |
|---|---|---|
| PTWAS | 0.621 | 0.038 |
| Single-variant | 0.584 | 0.021 |
| Elastic net | 0.634 | 0.031 |

Table S3: **Simulation Multi-INTACT results across a variety of expression and protein prediction models** For each simulated gene, protein and expression levels are predicted using the model indicated in the left-most column. The underlying causal DAG has $E \to Y$, $P \to Y$, and $P \to E$ effects. Realized power and false discovery rate (FDR) at the 5% control level are provided in the right-most columns.

| Multi-INTACT $\chi^2$ statistic formula | Multi-INTACT power | Multi-INTACT FDR |
|---|---|---|
| Individual-level data | 0.576 | 0.045 |
| Summary-level data (DAP-G formula) | 0.573 | 0.046 |
| Summary-level data (MultiXcan formula) | 0.576 | 0.045 |

Table S4: **Simulation Multi-INTACT results using both individual-level and summary-level data** We compare the individual-level data approach for calculating the chi-square statistic to two summary-level data approaches adapted from previously introduced methodology. Realized power and false discovery rate (FDR) at the 5% control level are provided in the right-most columns. Full details for each approach included in Supplemental Methods.

### Supplemental Methods

### 1 Details of the Multi-INTACT model

#### 1.1 Computation of the gene probability of putative causality

The gene probability of putative causality introduced in Equation (7) can be computed analytically using Bayes' rule as:

$$\Pr(\gamma \neq 0 \text{ or } \delta \neq 0 \mid \text{ data }) = \frac{\text{BF}\pi f(p_{coloc,E}, p_{coloc,P})}{1 - \pi f(p_{coloc,E}, p_{coloc,P}) + \text{BF}\pi f(p_{coloc,E}, p_{coloc,P})}. \tag{S1}$$

.

##### 1.1.1 Computation of gene-level Bayes factors

To compute the Bayes factor, we consider a general test statistic for testing the null model, $M_0$, by fitting regression model (6) in a frequentist setting. The common choices of the test statistics are the $F$ statistic, the multivariate Wald statistic ($W$), and likelihood ratio statistic ($LR$). Noticeably, all these test statistics can be represented as monotonic increasing functions of the squared canonical correlation of interest, $R^2$ [1], i.e.,

$$\begin{aligned} F &= \frac{R^2/2}{(1-R^2)/(N-2)} \\ W &= N\frac{R^2}{1-R^2} \\ LR &= -N\log(1-R^2) \end{aligned} \tag{S2}$$

We first transform the test statistic to a corresponding $Z^2$ statistic, such that the two-sided $p$-value of the new $Z$-statistic remains identical to the original $p$-value. (Note that the function of transformed $Z^2$, $\frac{Z^2}{N+Z^2}$, corresponds to a shrinkage estimate of $R^2$.) We then compute a Bayes factor using Wakefield's approximation [2],

$$\text{BF} = \sqrt{\frac{1}{1+K}} \exp\left(\frac{Z^2}{2}\frac{K}{1+K}\right). \tag{S3}$$

Note that the function of the hyperparameter $K$, $\frac{K}{K+N}$, can be interpreted as the prior expected value of $R^2$ under the alternative models. In practice, we put a discrete uniform prior on a grid of $K$ values (or equivalently, the corresponding prior $R^2$ values), i.e., $\{K : 1, 2, 4, 8, 16\}$, and obtain the final Bayes factor by model averaging. By default, our implementation uses the multivariate Wald statistic because of its computational convenience to accommodate GWAS summary statistics. Additionally, in our simulation studies and real data analysis, we find that the difference in Bayes factor results due to alternative choices of input test statistics and/or the grid of $K$ values are negligible.

##### 1.1.2 Computation of the prior $\pi$

The parameter $\pi$ in the composite prior is derived by pooling the $p$-values of the test statistics (e.g., the $W$ statistics) for $M_0$ from all candidate genes. It is analogous to the default TWAS prior used in INTACT (see equation S5 of [3]). The estimation procedure is similar to what is used in the $q$-value procedure [4]:

$$\hat{\pi} = 1 - \frac{\sum_{j=1}^{m} I(p_j > \lambda)}{m(1-\lambda)} \tag{S4}$$

The $p_j$ denote $p$-values derived from the multivariate Wald test statistics for $m$ genes. The parameter $\lambda$ is set to 0.5 as in INTACT (see Supplemental Methods of [3] for a discussion of the logic behind this choice of $\lambda$ and the origin of the estimator in Equation S4). The resulting $\pi$ represents a lower bound of the proportion of signals in all candidates when colocalization evidence is ignored.

##### 1.1.3 Prior choices for Multi-INTACT

The function $f(p_{\text{coloc,E}}, p_{\text{coloc,P}})$, ranging from 0 to 1, modifies the exchangeable prior $\pi$ for individual target genes based on their colocalization evidence. We provide multiple options for the $f$ function, all of which shrink the overall prior probability towards 0 when modest gene-level colocalization is lacking. The function takes the general form

$$f(p_{\text{coloc,E}}, p_{\text{coloc,P}}) = \begin{cases} g(h(p_{\text{coloc,E}}, p_{\text{coloc,P}})) & \text{if } h(p_{\text{coloc,E}}, p_{\text{coloc,P}}) \geq t \\ 0 & \text{if } h(p_{\text{coloc,E}}, p_{\text{coloc,P}}) < t, \end{cases} \tag{S5}$$

The default implementation, which we use for analyzing simulated and real data, uses the identity function for $g$ and $h(p_{\text{coloc,E}}, p_{\text{coloc,P}}) = max(p_{\text{coloc,E}}, p_{\text{coloc,P}})$. The hard threshold $t$ is a pre-defined value and set to 0.05 by default.

To offer flexibility to the user, we include additional $g$ functions in the Multi-INTACT software:

**Step**:

$$g(x) = \mathbf{1}(x \geq t) = \begin{cases} 1 & \text{if } x \geq t \\ 0 & \text{otherwise} \end{cases}$$

**Expit**:

$$\begin{aligned} g(x) &= \text{expit}\left[(x - 0.5)/D\right] \mathbf{1}(x \geq t) \\ &= \frac{1}{1 + \exp\left(-\frac{x-0.5}{D}\right)} \mathbf{1}(x \geq t) \end{aligned}$$

**Hybrid-Linear-Expit**:

$$g(x) = \begin{cases} 0 & x < t \\ \text{expit}\left[(x - 0.5)/D\right] & t \leq x < 0.5 \\ x & x \geq 0.5 \end{cases}$$

The parameter $t$ is 0.5 by default for the step prior and 0.05 by default for all other prior functions. The parameter $D$ impacts the steepness of the expit prior curve.

Finally, the software provides one alternative option for the $h$ function:

$$h(p_{\text{coloc,E}}, p_{\text{coloc,P}}) = 1 - (1 - p_{\text{coloc,E}})(1 - p_{\text{coloc,P}}), \tag{S6}$$

Equation S6 generally produces less-conservative quantities than $h(p_{\text{coloc,E}}, p_{\text{coloc,P}}) = max(p_{\text{coloc,E}}, p_{\text{coloc,P}})$, particularly when all pairwise colocalization probabilities are moderate.

##### 1.2 Multi-INTACT EM algorithm

Here we detail the EM algorithm for estimating the parameters $h_E$, $h_P$, and $h_{E+P}$.

First, we compute Wakefield Bayes factors for the models $M_E$, $M_P$, and $M_{E+P}$ using the TWAS z-scores, PWAS z-scores, and multivariate Wald statistics, respectively. Given colocalization evidence for the $i$th gene $f_i(p_{\text{coloc,E}}, p_{\text{coloc,P}})$, we can write the prior model probabilities for the $i$th gene as:

$$P_i(M_E) = h_E \pi f_i(p_{\text{coloc,E}}, p_{\text{coloc,P}}), \tag{S7}$$

$$P_i(M_P) = h_P \pi f_i(p_{\text{coloc,E}}, p_{\text{coloc,P}}), \tag{S8}$$

$$P_i(M_{E+P}) = h_{E+P} \pi f_i(p_{\text{coloc,E}}, p_{\text{coloc,P}}), \tag{S9}$$

where $h_E + h_P + h_{E+P} = 1$. With this model formulation, we can estimate the exchangeable priors $h_E, h_P, h_{E+P}$ using an EM algorithm.

Provided colocalization evidence $f_i(p_{\text{coloc,E}}, p_{\text{coloc,P}})$ and Bayes factors $\text{BF}_{E,i}, \text{BF}_{P,i}, \text{BF}_{E+P,i}$, we first initiate the EM algorithm by setting

$$h_E^{(0)} = h_P^{(0)} = h_{E+P}^{(0)} = 1/3. \tag{S10}$$

We define $\gamma_{0,i}$, $\gamma_{E,i}$, $\gamma_{P,i}$, and $\gamma_{E+P,i}$ as missing indicator variables indicating the model pertaining to each gene.The complete data log likelihood can be written as

$$
\begin{aligned}
l(h_E, h_P, h_{E+P}) = \sum_{i=1}^{p}[log(\mathrm{BF}_{E,i}) + log(h_E\pi + log(f_i(p_{\mathrm{coloc,E}}, p_{\mathrm{coloc,P}})))] * \mathbf{1}(\gamma_{i,E} = 1) \\
+ \sum_{i=1}^{p}[log(\mathrm{BF}_{P,i}) + log(h_P\pi + log(f_i(p_{\mathrm{coloc,E}}, p_{\mathrm{coloc,P}})))] * \mathbf{1}(\gamma_{i,P} = 1) \\
+ \sum_{i=1}^{p}[log(\mathrm{BF}_{E+P,i}) + log(h_{E+P}\pi + log(f_i(p_{\mathrm{coloc,E}}, p_{\mathrm{coloc,P}})))] * \mathbf{1}(\gamma_{i,E+P} = 1) \\
+ \sum_{i=1}^{p} log[(1-\pi)f_i(p_{\mathrm{coloc,E}}, p_{\mathrm{coloc,P}}) + 1 - f_i(p_{\mathrm{coloc,E}}, p_{\mathrm{coloc,P}})] * \mathbf{1}(\gamma_{i,0} = 1)
\end{aligned}
\tag{S11}
$$

**E-step** Let $h_E^{(t)}$, $h_P^{(t)}$,$h_{E+P}^{(t)}$ denote estimates in the $t$th iteration of the EM algorithm.

Let $Q = \sum_{l\in\{E,P,E+P\}} h_l^{(t)}\pi f_i(p_{\mathrm{coloc,E}}, p_{\mathrm{coloc,P}})\mathrm{BF}_{i,l}$

$$
\begin{aligned}
Pr(\gamma_{i,l}|D, h_E^{(t)}, h_P^{(t)}, h_{E+P}^{(t)}) = \frac{h_l^{(t)}\pi f_i(p_{\mathrm{coloc,E}}, p_{\mathrm{coloc,P}})\mathrm{BF}_{i,l}}{[(1-\pi)f_i(p_{\mathrm{coloc,E}}, p_{\mathrm{coloc,P}}) + 1 - f_i(p_{\mathrm{coloc,E}}, p_{\mathrm{coloc,P}})] + Q} \\
Pr(\gamma_{i,0}|D, h_E^{(t)}, h_P^{(t)}, h_{E+P}^{(t)}) = \frac{[(1-\pi)f_i(p_{\mathrm{coloc,E}}, p_{\mathrm{coloc,P}}) + 1 - f_i(p_{\mathrm{coloc,E}}, p_{\mathrm{coloc,P}})]}{[(1-\pi)f_i(p_{\mathrm{coloc,E}}, p_{\mathrm{coloc,P}}) + 1 - f_i(p_{\mathrm{coloc,E}}, p_{\mathrm{coloc,P}})] + Q},
\end{aligned}
\tag{S12}
$$

for $l \in \{E, P, E+P\}$.

**M-step** We update the current estimates of $h_E$, $h_P$, and $h_{E+P}$ by finding

$\left(h_E^{(t+1)}, h_P^{(t+1)}, h_{E+P}^{(t+1)}\right) = argmax(\sum_{i=1}^{p}\sum_{l\in\{E,P,E+P\}} log[h_l\pi Pr(\gamma_{i,l} = 1|D, h_E, h_P, h_{E+P})]$
$+ \sum_{i=1}^{p} log[(1-\pi)f_i(p_{\mathrm{coloc,E}}, p_{\mathrm{coloc,P}}) + 1 - f_i(p_{\mathrm{coloc,E}}, p_{\mathrm{coloc,P}})]Pr(\gamma_{i,0} = 1|D, h_E, h_P, h_{E+P})),$

subject to $h_E + h_P + h_{E+P} = 1$.

Using the method of Lagrange multipliers, we maximize the following expression with respect to $h_E, h_P, h_{E+P}$:

$\sum_{i=1}^{p}\left[\sum_{l\in\{E,P,E+P\}} log(h_l\pi)Pr(\gamma_{i,l}|D, h_E^{(t)}, h_P^{(t)}, h_{E+P}^{(t)})\right] + \frac{f_i(p_{\mathrm{coloc,E}}, p_{\mathrm{coloc,P}})Pr(\gamma_{i,0}|D, h_E^{(t)}, h_P^{(t)}, h_{E+P}^{(t)})}{[1-\pi]f_i(p_{\mathrm{coloc,E}}, p_{\mathrm{coloc,P}}) + 1 - f_i(p_{\mathrm{coloc,E}}, p_{\mathrm{coloc,P}})}$
$+ \lambda(\sum_{l\in\{E,P,E+P\}} h_l - 1)$

When we set derivatives with respect to $h_E$, $h_P$, and $h_{E+P}$ to zero, we find

$\frac{\sum_{i=1}^{p} Pr(\gamma_{i,l}|D, h_E^{(t)}, h_P^{(t)}, h_{E+P}^{(t)})}{h_l\pi} - \lambda = 0.$

Thus,

$h_l = \frac{\sum_{i=1}^{p} Pr(\gamma_{i,l}|D, h_E^{(t)}, h_P^{(t)}, h_{E+P}^{(t)})}{\lambda\pi}$

$\lambda = \sum_{i=1}^{p} \sum_{l \in \{0,E,P,E+P\}} P(\gamma_{i,l}|D, h_E^{(t)}, h_P^{(t)}, h_{E+P}^{(t)}) = p$

Therefore,

$h_l^{(t+1)} = \frac{1}{p} \sum_{i=1}^{p} P(\gamma_{i,l}|D, h_E^{(t)}, h_P^{(t)}, h_{E+P}^{(t)}),\ l \in \{E, P, E+P\}$

#### 1.3 Computation with GWAS summary statistics

When only summary-level data is available, we can approximate a $\chi^2$ statistic from the regression in Equation (6). We emphasize that although this approximation is convenient for practical use, we strongly recommend using individual-level data whenever possible. In this section, we describe two approximation procedures which make different assumptions about the underlying model.

**Approach 1:**

The first approach that we describe requires that GWAS summary statistics (single-SNP z scores), molecular phenotype prediction weights, and an appropriate LD reference panel for the GWAS SNPs are available.

The estimated effect size vector of interest $\hat{\beta} = [\hat{\gamma}, \hat{\delta}]$ can be expressed as a function of the genotype matrix, prediction weight matrix $W$, and complex trait vector:

$$\hat{\beta} = [X^T X]^{-1} X^T Y = [(GW)^T GW]^{-1} (GW)^T Y = [W^T G^T GW]^{-1} W^T G^T Y$$

and is is asymptotically normal:

$$\hat{\beta} \sim N(\beta, Var(\hat{\beta}))$$

, where $Var(\hat{\beta}) = (X^T X)^{-1} \sigma_Y^2$.

The individual-level test statistic corresponding to the hypothesis test (Equation **??**) is

$$\hat{\beta}^T [Var(\hat{\beta})]^{-1} \hat{\beta}$$

$$= Y^T GW[W^T G^T GW]^{-1}[W^T G^T GW]\sigma_Y^{-2}[W^T G^T GW]^{-1} W^T G^T Y$$

$$= Y^T GW \sigma_Y^{-2}[W^T G^T GW]^{-1} W^T G^T Y$$

Using the fact that $G^T G = \Lambda R \Lambda$ and $G^T Y = \sigma_g \Lambda \mathbf{z}$ [5], where $R$ is the SNP correlation matrix, $\Lambda = diag\left[\sqrt{\mathbf{g_1^T g_1}}, ..., \sqrt{\mathbf{g_p^T g_p}}\right]$, $\mathbf{z}$ is a vector of single-SNP GWAS z scores, and $\sigma_g$ is the true residual error standard deviation across all single-variant GWAS regression models (assumed to be constant across variants), we can express $\hat{\beta}$ as

$$\mathbf{z}^T \Lambda W \frac{\sigma_g}{\sigma_Y}[W^T \Lambda R \Lambda W]^{-1} \frac{\sigma_g}{\sigma_Y} W^T \Lambda \mathbf{z}$$

.

Furthermore, assuming that $\sigma_g \approx \sigma_Y$, we find that the test statistic can be approximated as

$$\mathbf{z}^T \Lambda W[W^T \Lambda R \Lambda W]^{-1} W^T \Lambda \mathbf{z}$$

.

The GWAS summary statistics $\mathbf{z}$ are commonly provided, while prediction weights $W$ can be found from databases such as PredictDB [6, 7, 8]. We assume the matrices $R$ and $\Lambda$ can be estimated from an appropriate LD reference panel. Following the recommendation of previous work [9], we advise only including variants that were used in each molecular trait prediction model to generate this test statistic.

**Approach 2:**

The second approach we describe can directly work with marginal z scores derived from a TWAS method. In brief, we approximate the $\chi^2$ test statistic using the MultiXcan framework [9], where we consider multiple predicted molecular phenotype gene products rather than predicted gene expression in multiple tissues. This approach assumes that the residual error variance from the marginal TWAS regressions is identical to that from the multiple regression of the complex trait on all predicted molecular phenotypes. The approximation of the test statistic takes the form

$$\mathbf{z_{TWAS}^T}[\mathbf{Cor}(\mathbf{X})]^{-\mathbf{1}}\mathbf{z_{TWAS}}$$

,

where $\mathbf{z_{TWAS}}$ is a vector of marginal TWAS z scores (of length 2 if considering gene expression and protein levels), and $\mathbf{X}$ is a standardized predicted molecular phenotype matrix (each column has mean 0 and standard deviation 1). In practice, $\mathbf{Cor}(\mathbf{X})$ can be approximated using a genotype covariance matrix from an appropriate reference panel as well as the TWAS weights $W$. For the full derivation of the approximation, see [9]. This approach is convenient compared to the first approach if one already has access to marginal TWAS association results or the matrix $\Lambda$ cannot be estimated from a reference panel.

**A comparison of Approaches 1 & 2:**

Although approaches 1 and 2 represent valid approximations of the joint regression test statistic, they each rely on different assumptions and thus are not the same. To illustrate the accuracy of each approximation compared to the individual-level data approach, using one of our simulated data sets, we plot the gene-level test statistics from each approximation against the test statistic using individual-level data (Figure S7). We observe that the test statistic approximations both Approaches 1 and 2 are generally quite accurate in approximating the individual-level statistic, with a drop in accuracy at high individual-level statistic values. This is not a concern since this will generally not affect the power of the analysis.

#### 2 Design of additional simulations

We simulate 9 separate data sets, each representing one of the DAGs from Table S1. Each subsection describes the simulation algorithm pertaining to one data set corresponding to one of the Table S1 entries.

##### 2.1 $E \to Y$, $P \to Y$, and $P \to E$

1. For the $i$th gene, randomly select 2 causal cis-eQTLs ($g_{i,1}$, $g_{i,2}$) and 2 cis-pQTLs, ($g_{i,2}$, $g_{i,3}$), with exactly one overlapping causal QTL ($g_{i,2}$). Randomly select a distinct causal GWAS SNP ($g_{i,4}$). Draw a causality indicator variable $\eta_i$ from a Bernoulli(0.2) distribution. Beyond this point, we omit the $i$ subscript, but note that all parameters and variables are gene-specific.

2. Simulate protein data as

$$P = g_2\beta_{2,P} + g_3\beta_{3,P} + e_P,$$

where $\beta_{2,P}$ and $\beta_{3,P}$ are drawn independently from a $N(0, 0.6^2)$ distribution and $e_P$ are drawn independently from a $N(0, 1)$ distribution.

3. Simulate expression data as

$$E = g_1\beta_{1,E} + g_2\beta_{2,E} + P\delta_E + e_E,$$

where $\beta_{1,E}$, $\beta_{2,E}$, and $\delta_E$ are drawn from a $N(0, 0.6^2)$ distribution and $e_E$ are drawn independently from a $N(0, 1)$ distribution.

4. Simulate complex trait data.

If $\eta = 1$,

$$Y = E\gamma_Y + P\delta_Y + g_4\beta_4 + e_Y,$$

where $\gamma_Y$, $\delta_Y$, and $\beta_4$ are drawn from a $N(0, 0.6^2)$ distribution and $e_Y$ are drawn independently from a $N(0, 1)$ distribution.

If $\eta = 0$,

$$Y = g_4\beta_4 + e_Y$$

.

#### 2.2 $P \to Y$ and $P \to E$

1. Follow steps 1-3 of section 2.1.

2. Simulate complex trait data.

   If $\eta = 1$,

   $$Y = P\delta_Y + g_4\beta_4 + e_Y,$$

   where $\delta_Y$, and $\beta_4$ are drawn from a $N(0, 0.6^2)$ distribution and $e_Y$ are drawn independently from a $N(0, 1)$ distribution.

   If $\eta = 0$,

   $$Y = g_4\beta_4 + e_Y$$

   .

#### 2.3 $E \to Y$ and $P \to E$

1. Follow steps 1-3 of section 2.1.

2. Simulate complex trait data.

   If $\eta = 1$,

   $$Y = E\gamma_Y + g_4\beta_4 + e_Y,$$

   where $\gamma_Y$, and $\beta_4$ are drawn from a $N(0, 0.6^2)$ distribution and $e_Y$ are drawn independently from a $N(0, 1)$ distribution.

   If $\eta = 0$,

   $$Y = g_4\beta_4 + e_Y$$

   .

#### 2.4 $E \to Y$ and $P \to Y$ (no effects between $E$ and $P$)

1. Follow steps 1-2 of section 2.1.
2. Simulate expression data as

$$E = g_1\beta_{1,E} + g_2\beta_{2,E} + e_E,$$

where $\beta_{1,E}$ and $\beta_{2,E}$ are drawn from a $N(0, 0.6^2)$ distribution and $e_E$ are drawn independently from a $N(0, 1)$ distribution.

3. Follow step 4 of section 2.1.

#### 2.5 $P \to Y$ (no effects between $E$ and $P$)

1. Follow steps 1-2 of section 2.4.
2. Follow step 2 section 2.2.

#### 2.6 $E \to Y$ (no effects between $E$ and $P$)

1. Follow steps 1-2 of section 2.4.
2. Follow step 2 section 2.3.

#### 2.7 $E \to Y$, $P \to Y$, and $E \to P$

1. Follow step 1 of section 2.1.
2. Follow step 2 from section 2.4.
3. Simulate protein data as

$$P = g_2\beta_{2,P} + g_3\beta_{3,P} + E\gamma_P + e_P,$$

where $\beta_{2,P}$, $\beta_{3,P}$, and $E\gamma_P$ are drawn independently from a $N(0, 0.6^2)$ distribution and $e_P$ are drawn independently from a $N(0, 1)$ distribution.

4. Follow step 4 of section 2.1

#### 2.8 $P \rightarrow Y$ and $E \rightarrow P$

1. Follow steps 1-3 of section 2.7.
2. Follow step 2 of section 2.2

#### 2.9 $E \rightarrow Y$ and $E \rightarrow P$

1. Follow steps 1-3 of section 2.7.
2. Follow step 2 of section 2.3

### 3 METSIM Metabolon Metabolite GWAS data

The METSIM study comprises 10,197 men in Kuopio, Finland. Participants aged 45 to 74 were examined in baseline visits from 2005 to 2010. Those who were non-Finnish (n=21), failed whole-genome sequencing (WGS) (n=65), had sex mismatch(n=3), and/or lacked body mass index measurements (n=1) were excluded. Metabolon, Inc (Durham, North Carolina, USA)[10] performed non-targeted metabolomics profiling on EDTA-plasma samples. Samples were obtained after ≥10-hour overnight fasts during baseline visits. First, methanol extraction of biochemicals was applied; then, non-targeted relative quantitative liquid chromatography–tandem mass spectrometry Metabolon DiscoveryHD4 platform was applied to assay 1,544 metabolites. A randomized batch design was used, where batches contained ∼144 METSIM samples and 20 well-characterized human-EDTA plasma samples for quality control. Data processing, including peak quantification and data scaling, was performed for all 10.188 samples together. We used area under the curve to quantify raw mass spectrometry peaks for each metabolite. Overall process variablility was evaluated by the median relative standard deviation for endogenous metabolites that were present in all 20 technical replicates in each batch. In order to adjust for variation caused by day-to-day instrument tuning differences and columns used for biochmical extraction, we scaled the raw peak quantification to the median for each metabolite by batch.

Illumina HiSeq X Ten instruments were used for WGS, targeting a mean depth of at least 30x (paired-end, 150 bp reads). PCR-free library preparation kits from KAPA Biosystems were used for all sequencing. To process samples, CBCL files were converted to FASTQ-formatted reads. Reads were assigned to samples using bcl2fastq conversion software (Illumina Inc., San Diego, CA). Sample-specific FASTQ files were aligned to the GRCh38 genome reference with BWA-mem. Aligned reads were evaluated in BAM files. The Picard MarkDuplicates tool was usied to identify and flag duplicate reads. GVCF files for each individual sample The WeCall variant caller was used to produce GVCF files for each individual sample, which identified SNVs and INDELs as compared to the reference. The samtools tool was used to convert BAM files to CRAM files.

The following criteria were used for quality control: sex discrepancy between genetically-determined and self-reported sex, high rate of heterozygosity/contamination, low sequencing coverage, geneticallyidentified duplicates, and discordance between whole-exome sequencing and array sequencing.

### Supplemental References

### References

[1] Magee, L. R 2 measures based on wald and likelihood ratio joint significance tests. *The American Statistician* **44**, 250–253 (1990).

[2] Wakefield, J. Bayes factors for genome-wide association studies: comparison with p-values. *Genetic Epidemiology: The Official Publication of the International Genetic Epidemiology Society* **33**, 79–86 (2009).

[3] Okamoto, J. *et al.* Probabilistic integration of transcriptome-wide association studies and colocalization analysis identifies key molecular pathways of complex traits. *The American Journal of Human Genetics* **110**, 44–57 (2023).

[4] Storey, J. D. & Tibshirani, R. Statistical significance for genomewide studies. *Proceedings of the National Academy of Sciences* **100**, 9440–9445 (2003).

[5] Lee, Y., Luca, F., Pique-Regi, R. & Wen, X. Bayesian multi-snp genetic association analysis: control of fdr and use of summary statistics. *BioRxiv* 316471 (2018).

[6] Gamazon, E. R. *et al.* A gene-based association method for mapping traits using reference transcriptome data. *Nature genetics* **47**, 1091–1098 (2015).

[7] Barbeira, A. N. *et al.* Exploring the phenotypic consequences of tissue specific gene expression variation inferred from gwas summary statistics. *Nature communications* **9**, 1825 (2018).

[8] Barbeira, A. N. *et al.* Exploiting the gtex resources to decipher the mechanisms at gwas loci. *Genome biology* **22**, 1–24 (2021).

[9] Barbeira, A. N. *et al.* Integrating predicted transcriptome from multiple tissues improves association detection. *PLoS genetics* **15**, e1007889 (2019).

[10] Evans, A. M. *et al.* High resolution mass spectrometry improves data quantity and quality as compared to unit mass resolution mass spectrometry in high-throughput profiling metabolomics. *Metabolomics* **4**, 1 (2014).
